# Several geranylgeranyl diphosphate synthase isoforms supply metabolic substrates for carotenoid biosynthesis in tomato

**DOI:** 10.1101/2020.07.08.194076

**Authors:** M. Victoria Barja, Miguel Ezquerro, Gianfranco Diretto, Igor Florez-Sarasa, Elisenda Feixes, Alessia Fiore, Rumyana Karlova, Alisdair R. Fernie, Jules Beekwilder, Manuel Rodriguez-Concepcion

**Affiliations:** Centre for Research in Agricultural Genomics (CRAG) CSIC-IRTA-UAB-UB, Campus UAB Bellaterra, 08193 Barcelona, Spain; Italian National Agency for New Technologies, Energy, and Sustainable Development, Casaccia Research Centre, 00123 Rome, Italy; Laboratory of Plant Physiology, Wageningen University and Research, 6700AA Wageningen, The Netherlands; Max-Planck-Institut für Molekulare Pflanzenphysiologie, 14476 Potsdam-Golm, Germany; BU Bioscience, Wageningen University and Research, 6700AA Wageningen, The Netherlands; Institute for Plant Molecular and Cell Biology (IBMCP), CSIC-Universitat Politècnica de València, 46022 Valencia, Spain

## Abstract

Geranylgeranyl diphosphate (GGPP) produced by GGPP synthase (GGPPS) serves as a precursor for many plastidial isoprenoids, including carotenoids. Here we show that five different GGPPS isoforms exist in tomato (*Solanum lycopersicum*). From these, SlGGPPS1, 2 and 3 (or SlG1-3 in short) produce GGPP in plastids and exhibit similar kinetic parameters. Phytoene synthase (PSY) converts GGPP into phytoene, the first committed intermediate of the carotenoid pathway. Gene expression and co-expression network analyses showed a preferential association of individual GGPPS and PSY isoforms in processes linked to carotenoid biosynthesis such as root mycorrhization, seedling deetiolation and fruit ripening. Co-immunoprecipitation experiments showed that SlG2, but not SlG3, physically interacts with PSY proteins. By contrast, CRISPR-Cas9 mutants defective in SlG3 showed a stronger impact on carotenoid levels and derived metabolic, physiological and developmental phenotypes that those impaired in SlG2. Double mutants with a simultaneous knockout of both genes could not be found. Our work demonstrates that the bulk of GGPP production in tomato chloroplasts and chromoplasts relies on two cooperating GGPPS paralogs, unlike other plant species such as *Arabidopsis thaliana*, rice or pepper, which produce their essential plastidial isoprenoids using a single GGPPS isoform.

## INTRODUCTION

Isoprenoids are essential biological molecules in all living organisms. In particular, plants are the main source of the enormous structural and functional variety that characterizes this family of compounds (Pulido et al., 2012; Tholl, 2015). The building blocks for the biosynthesis of all isoprenoids are isopentenyl diphosphate (IPP) and its double-bond isomer dimethylallyl diphosphate (DMAPP). These five-carbon (C5) universal isoprenoid units are produced in plants by the mevalonic acid (MVA) pathway in the cytosol and the methylerythritol 4-phosphate (MEP) pathway in plastids (Vranová et al., 2013; Rodríguez-Concepción and Boronat, 2015). Short-chain prenyltransferases (SC-PTs) subsequently condense one or more molecules of IPP to one molecule of DMAPP giving rise to C10, C15, C20 and C25 prenyl diphosphates known as geranyl diphosphate (GPP), farnesyl diphosphate (FPP), geranylgeranyl diphosphate (GGPP), and geranylfarnesyl diphosphate (GFPP), respectively. These molecules are the immediate precursors for downstream pathways leading to the production of the main groups of isoprenoids. GPP is produced in plastids as the precursor of C10 monoterpenes. FPP is mainly produced in the cytosol and used to synthesize C15 sesquiterpenes and C30 triterpenes (including phytosterols). GGPP is mostly produced in plastids, serving as the precursor of gibberellins and photosynthesis-related isoprenoids such as chlorophylls, carotenoids, tocopherols, plastoquinone and phylloquinones. However, GGPP is also used for the production of C20 diterpenes in the cytosol, and both FPP and GGPP are produced in mitochondria for ubiquinone and diterpenoid biosynthesis. GFPP is used to produce C25 sesterterpenes in different cell compartments. SC-PTs are encoded by gene families in most plants and they are typically found in different cell compartments, consistent with the requirement of their specific prenyl diphosphate products in different subcellular locations. Prediction of specific products and cell targeting based solely on their protein sequences is still a challenge, making experimental evidence necessary to ascertain their biological role (Cunillera et al., 1996, 1997; Gaffe et al., 2000; Beck et al., 2013; Jones et al., 2013; Nagel et al., 2015; Wang et al., 2016a; Zhou et al., 2017; Zhou and Pichersky, 2020).

Carotenoids are one of the most studied groups of plant isoprenoids. These C40 tetraterpenes are greatly demanded by cosmetic and agro-food industries as natural red to yellow pigments and provide benefits for human health, *e.g.* as precursors of vitamin A and other biologically active molecules (Sandmann, 2015; Rodriguez-Concepcion et al., 2018). In plants, carotenoids have different functions. In photosynthetic tissues, they are required for the assembly of the photosynthetic apparatus, contribute to light harvesting and are essential for photoprotection by dissipating excess light energy as heat and by scavenging reactive oxygen species. They are also fundamental in growth regulation, since they are the precursors of retrograde signals and phytohormones such as abscisic acid (ABA) and strigolactones. As a secondary role, carotenoids provide distinctive colors to flowers and fruits to attract pollinators and seed dispersal animals (Nisar et al., 2015; Yuan et al., 2015). In plants, carotenoids are produced and stored in plastids, including chloroplasts and chromoplasts (Ruiz-Sola and Rodríguez-Concepción, 2012; Sun et al., 2018). MEP-derived IPP and DMAPP are converted into GGPP by plastidial GGPP synthase (GGPPS) isoforms and then GGPP is transformed into phytoene by phytoene synthase (PSY) enzymes. The production of phytoene, the first committed intermediate of the carotenoid pathway, is considered to be a major rate-determining step regulating the metabolic flux through this pathway (Fraser et al., 2002). In tomato (*Solanum lycopersicum*), three PSY-encoding genes control carotenoid biosynthesis in different tissues. *PSY1* expression is boosted during ripening to produce carotenoids involved in the pigmentation of the fruit (Bartley et al., 1992; Fray and Grierson, 1993; Giorio et al., 2008; Kachanovsky et al., 2012). *PSY2* is expressed in all tissues, including fruits, but transcript levels are much higher than those of *PSY1* in photosynthetic tissues, where carotenoids are required for photosynthesis and photoprotection (Bartley and Scolnik, 1993; Giorio et al., 2008). Lastly, *PSY3* is mainly expressed in roots and it is induced during mycorrhization (Walter et al., 2015; Stauder et al., 2018), when carotenoid biosynthesis is up-regulated to produce strigolactones and apocarotenoid molecules essential for the establishment of the symbiosis (Fester et al., 2002, 2005; Baslam et al., 2013; Ruiz-Lozano et al., 2016; Stauder et al., 2018). Whether the corresponding PSY isoforms use GGPP supplied by different GGPPS isoforms remains unknown.

Several GGPP synthase (GGPPS) paralogs have been retained in plants during evolution (Beck et al., 2013; Zhang et al., 2015; Ruiz-Sola et al., 2016a, 2016b; Zhou et al., 2017; Wang et al., 2018). However, a single GGPPS isoform appears to produce the GGPP substrate needed for the production of carotenoids and other plastidial isoprenoids in the three plant species whose GGPPS families have been best characterized to date: *Arabidopsis thaliana,* rice (*Oryza sativa*) and pepper (*Capsicum annuum*) (Ruiz-Sola et al., 2016a, 2016b; Zhou et al., 2017; Wang et al., 2018). While tomato has become one of the best plant systems to study the biosynthesis of carotenoids and its regulation, we still have a very incomplete picture of the GGPPS family in this plant. In particular, the GGPPS isoform(s) required for the production of carotenoids in photosynthetic tissues (*e.g.* for photoprotection), fruits (*e.g.* for pigmentation) or roots (*e.g.* for mycorrhization) remain unknown. Here we identify the plastidial GGPPS set in tomato and provide clues to understand how the supply of plastidial GGPP for the synthesis of carotenoids with different biological functions in particular tissues is regulated in this important crop plant.

## RESULTS

### The tomato GGPPS protein family includes three plastidial isoforms

Several genes encoding proteins with homology to GGPPS enzymes can be found in the tomato genome (Table S1). Two of them were previously reported and named *GGPPS1* (*Solyc11g011240*) and *GGPPS2* (*Solyc04g079960*) (Ament et al., 2006; Fraser et al., 2007; Stauder et al., 2018; Zhou and Pichersky, 2020). Besides them, herein referred to as *SlG1* and *SlG2*, three more putative GGPPS paralogs were found using as a query the *Arabidopsis thaliana GGPPS11* (*AtG11*) gene (At4g36810). We named them *SlG3* (*Solyc02g085700*), *SlG4* (*Solyc02g085710*), and *SlG5* (*Solyc02g085720*). In our phylogenetic analyses (Figure 1), SlG1 was found to cluster with *Arabidopsis* GGPPS1 (AtG1), a mitochondrial isoform found to produce both GGPP and GFPP, whereas SlG2 and SlG3 were more closely related to GGPPS2 (AtG2) and GGPPS11 (AtG11), the two plastid-targeted *Arabidopsis* isoforms that mainly produce GGPP (Nagel et al., 2015; Wang et al., 2016a). SlG4 and SlG5 grouped in a separated cluster (Figure 1). All five tomato GGPPS candidates harbored the two essential catalytic motifs required for prenyltransferase activity (FARM and SARM) and a methionine (M) residue in the fifth position upstream to the FARM shown to be essential for GGPP production (Figure 1) (Figure S1) (Nagel et al., 2015; Wang et al., 2016a).

**Figure 1.**
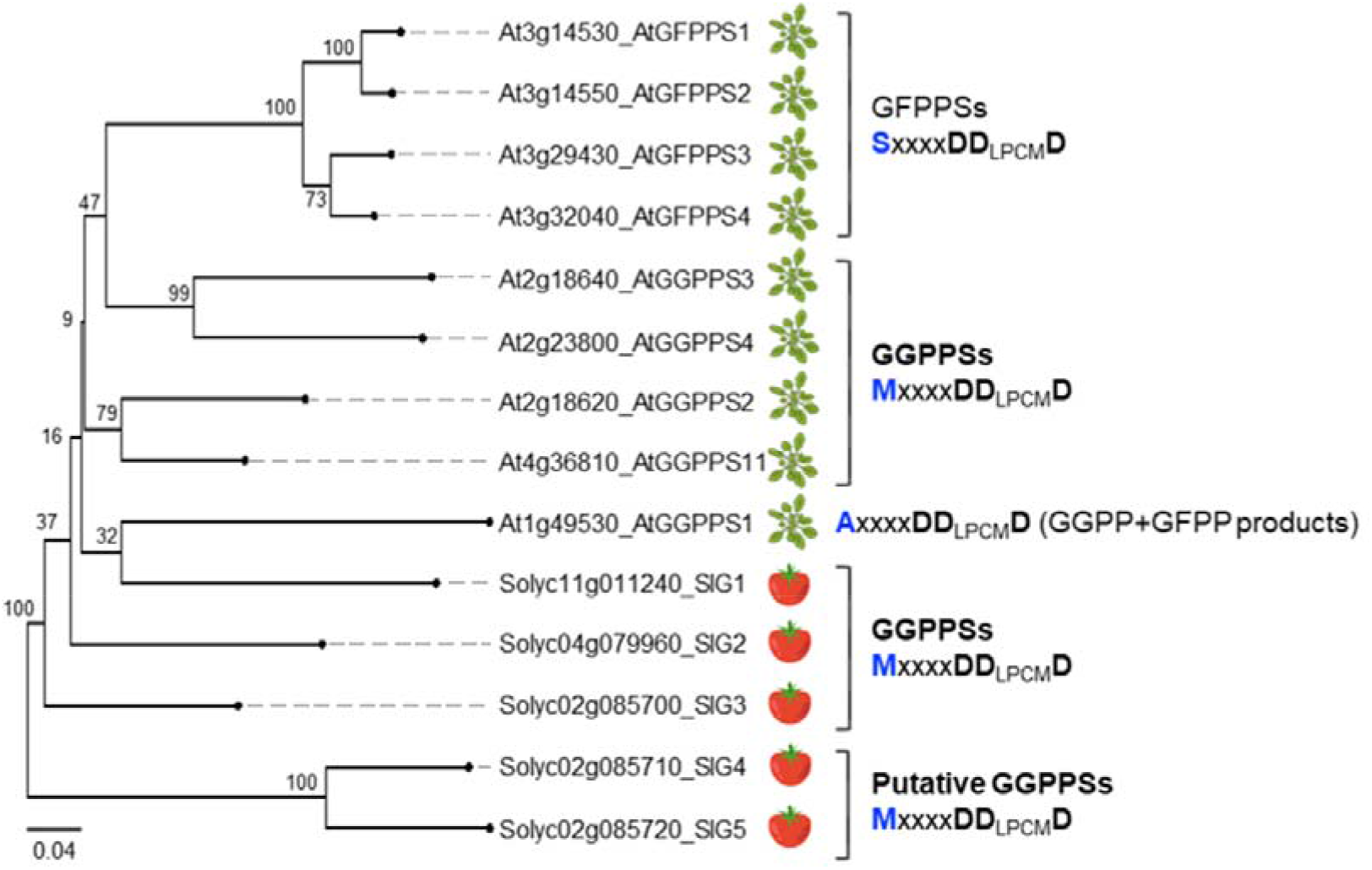
Phylogenetic tree of GGPP and GFPP enzymes. Arabidopsis and tomato protein sequences lacking their predicted transit peptides were used to generate an unrooted Neighbor-Joining tree. The percentage of trees in which the associated sequences clustered together is indicated in each branch. The scale bar represents the mean number of substitutions per site. The chain-length determination (CLD) region (*i.e.* the FARM and the previous five residues) is indicated for each enzyme lineage. Residues determining the product length are shown in blue and aspartate (D) residues are shown in black; ‘x’ represents any amino acid. See Table S1 for accessions.

Next, we carried out experiments to ascertain the subcellular localization and enzymatic parameters of the identified GGPPS candidates, SlG1-5 (Figure 2). First, we fused their complete sequences to a C-terminal green fluorescent protein (GFP) reporter and cloned the resulting constructs under the control of the *35S* promoter. Following agroinfiltration of tobacco (*Nicotiana benthamiana*) leaves, the green fluorescence emitted by each chimeric protein was analysed by confocal microscopy at 3 dpi (days post-infiltration) (Figure 2A). The GFP fusions of SlG1-3 showed a clear chloroplast localization as their GFP fluorescence overlapped with chlorophyll autofluorescence. By contrast, the signal corresponding to the SlG5-GFP fusion protein was detected as a punctate pattern typical of mitochondrial proteins (Figure 2A). This subcellular compartmentalization is in agreement with *in silico* predictions by *TargetP* and *ChloroP* algorithms (Emanuelsson et al., 2007). Fusions of the N-terminal 120 aa sequence of the tomato GGPPS candidates to GFP were also reported to localize in chloroplasts (SlG1/GGPPS1, SlG2/GGPPS2 and SlG3/GGPPS3) or mitochondria (SlG5/TPT2) when expressed in Arabidopsis protoplasts (Zhou and Pichersky, 2020). In the case of SlG4, the indicated algorithms predicted a plastidial targeting but we observed a predominantly cytosolic localization (Figure 2A) with only some cells occasionally exhibiting a plastidial localization of the SlG4-GFP reporter (Figure S2). Strikingly, a mitochondrial pattern was observed in Arabidopsis protoplasts using an N-terminal fragment of the SlG4/TPT1 protein fused to GFP (Zhou and Pichersky, 2020). These data, together with the observation that only SlG1, SlG2 and SlG3 produce GGPP *in vitro* (Zhou and Pichersky, 2020), led us to discard SlG4 and SlG5 and focus on the three tomato GGPPS enzymes with a clear plastidial location for further experiments.

**Figure 2.**
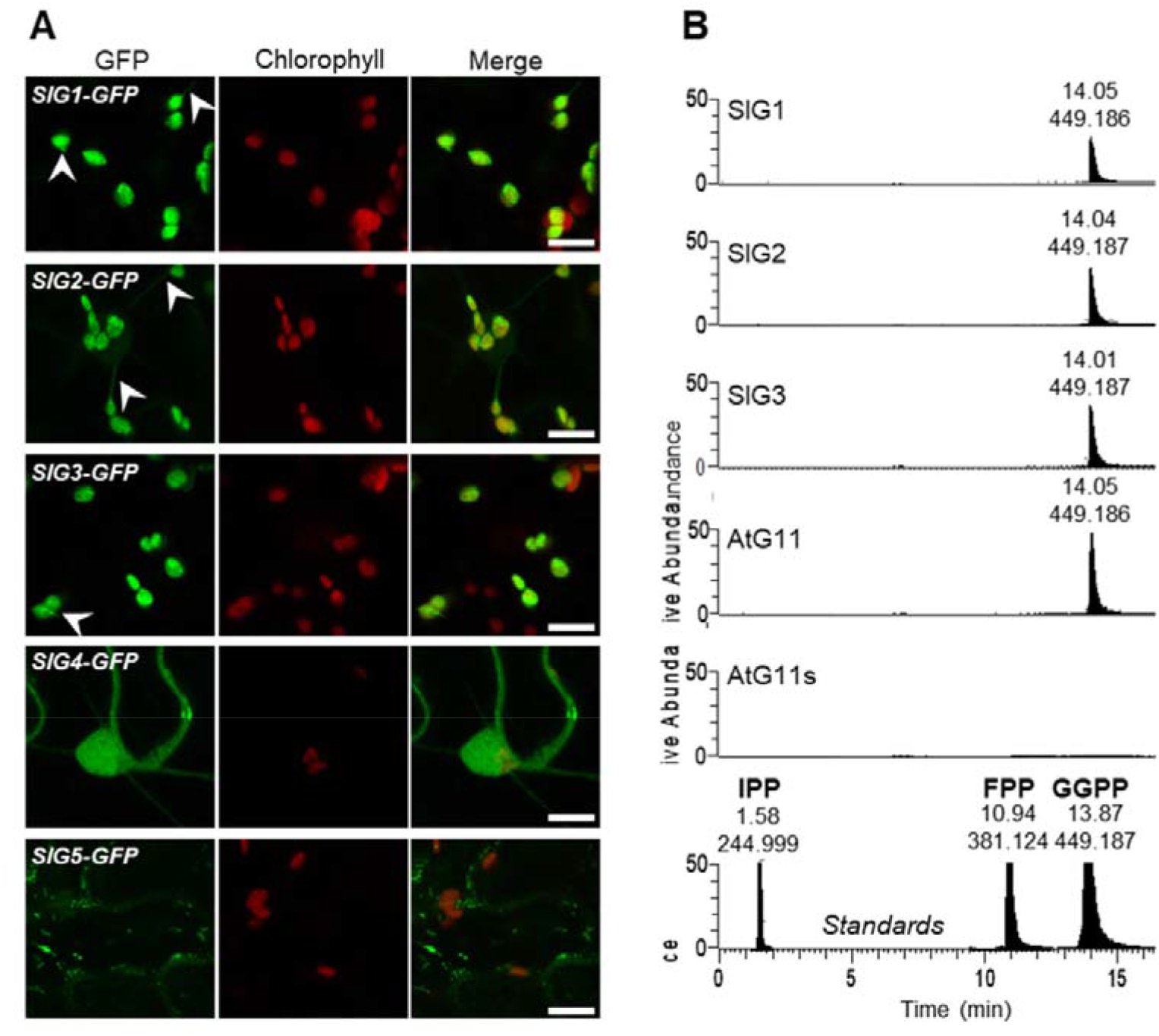
Subcellular localization and activity of tomato GGPPS candidates. **(A)** Representative confocal microscopy images of *N. benthamiana* leaf cells transiently expressing the indicated GFP fusion proteins. For each construct, GFP fluorescence (green, first column), chlorophyll autofluorescence (red, second column) and merged images of the same field (green and red, third column) are shown. White arrowheads mark stromules. Bars, 10 μm. **(B)** LC-MS chromatograms of reaction products. Extracts of *E. coli* cells overproducing the indicated recombinant proteins (with an N-terminal 6x-His tag instead of their predicted plastid-targeting peptide) were incubated with IPP and DMAPP and the products were analyzed by LC-MS. Prenyl diphosphates were detected using mass-to-charge (m/z) ratios: 244.998 (IPP), 313.061 (GPP), 381.123 (FPP), 449.186 (GGPP) and 518.254 (GFPP). Retention times of available standards is also shown.

### SlG1, SlG2 and SlG3 are GGPP-producing enzymes with similar kinetic properties

To investigate whether SlG1-3 proteins produce GGPP with similar efficiency, we set up *in vitro* activity assays followed by the analysis of the reaction products by LC-MS (Figure 2B). For expression in *Escherichia coli* cells, the three tomato isoforms were separately cloned without their predicted plastid-targeting sequences to aid solubility and fused to a N-terminal 6x-histidine tag to facilitate purification (Figure S3). As positive and negative controls, we used the Arabidopsis AtG11 (active) and AtG11s (inactive) proteins (Ruiz-Sola et al., 2016b). *E. coli* transformants were induced to produce the corresponding recombinant proteins (Figure S3) and whole-cell protein extracts were directly used for activity assays in the presence of IPP and DMAPP. LC-MS analysis of the reaction products showed that SlG1, SlG2, SlG3 and AtG11 (but no AtG11s) produced only GGPP (Figure 2B). Other prenyl diphosphate such as GPP, FPP or GFPP were not detected in the assays. These results confirmed that these three tomato proteins are true GGPPS enzymes, in agreement with recently reported data (Zhou and Pichersky, 2020).

In order to analyze the kinetic properties of the identified tomato GGPPS isoforms, we then purified the recombinant proteins taking advantage of their 6x-histidine tag, using again AtG11 as a positive control (Figure S3). Enzymatic assays with the purified enzymes (Barja and Rodríguez-Concepción, 2020) showed that all tested GGPPS proteins exhibited a similar optimal pH around 7.5 (Figure S3), as expected for stromal enzymes (Höhner et al., 2016). The kinetic parameters Km (an estimator of the apparent affinity for the IPP and DMAPP substrates) and Vmax exhibited very similar values among the three tomato enzymes (Table 1). They are also similar to those obtained for AtG11 here and elsewhere (Wang and Dixon, 2009; Camagna et al., 2019). We can therefore conclude that tomato SlG1, SlG2 and SlG3 and Arabidopsis AtG11 are true plastidial GGPPS enzymes with very similar kinetic properties.

**Table 1.** Kinetic parameters of plastidial GGPPS enzymes. Values correspond to the mean±SD of three independent experimental replicates (n=3).

| | DMAPP<br>(+100 μM IPP) | | IPP<br>(+100 μM DMAPP) | |
| --- | --- | --- | --- | --- |
| | Km<br>(μM) | Vmax<br>(nmol·min$^{-1}$ ·mg$^{-1}$) | Km<br>(μM) | Vmax<br>(nmol·min$^{-1}$ ·mg$^{-1}$) |
| SIG1 | 31.82 ± 2.92 | 47.47 ± 1.40 | 74.18 ± 7.55 | 59.87 ± 2.73 |
| SIG2 | 49.55 ± 5.31 | 38.87 ± 1.53 | 79.75 ± 8.33 | 36.73 ± 1.73 |
| SIG3 | 45.75 ± 6.81 | 26.13 ± 1.40 | 45.92 ± 4.86 | 29.13 ± 1.13 |
| AtG11 | 32.86 ± 4.86 | 21.53 ± 1.07 | 38.49 ± 4.94 | 24.13 ± 1.07 |

### Gene expression profiles suggest a major role of SlG2 and SlG3 in chloroplasts and chromoplasts

Analysis of public gene expression databases showed that the genes encoding SlG1-3 enzymes were expressed in roots, leaves and flowers (Figure S4). Of these, the most highly expressed gene is *SlG3* followed by *SlG2*, while *SlG1* transcripts are present at very low levels. *SlG2* and *SlG3*, but not *SlG1,* are also expressed at high levels in fruit pericarp and seed tissues (Figure S4). As an initial approach to get an insight on the possible functions of these individual isoforms, we performed a gene co-expression network (GCN) analysis. This is a powerful tool to infer biological functions by applying the simple concept of “guilt-by-association”, which means that a gene significantly associated with a molecular pathway at transcriptional level may share similar functions (Oliver, 2000). Because we were interested in understanding the contribution of SlG1, SlG2 and SlG3 to the production of carotenoids and related isoprenoids in plastids, we performed targeted GCN analyses with GGPP-related plastidial isoprenoid biosynthetic pathways in different tomato tissues (Figure 3) (Figure S5). By using publicly available databases for plant comparative genomics (*PLAZA 4.0*, *Phytozome*), we searched for tomato homologs of the pathways that supply GGPPS substrates (MEP pathway) and consume GGPP to produce carotenoids, chlorophylls, tocopherols, phylloquinone, plastoquinone, gibberellins, strigolactones and ABA (Table S2). Then, we retrieved their expression data from *TomExpress* (Zouine et al., 2017) in vegetative (i.e. photosynthetic) and fruit tissues and calculated their correlation with those of *SlG1, SlG2* and *SlG3* using pairwise Pearson correlations. It was not possible to obtain correlation data for tomato roots since only two expression values are available for each gene in the *TomExpress* database. GCN analyses are shown in Figure 3 and Figure S5, and positive significant correlations (ρ≥0.55) are listed in Table S3. The GCN analysis showed that *SlG1* is poorly co-expressed with the query genes (Figure 3). By contrast, *SlG2* and, to a lower extent, *SlG3* are highly connected to plastidial isoprenoid biosynthetic genes in green tissues. Connectivity was lower in fruit, and in this case it was a bit higher for *SlG3* (Figure 3). Remarkably, the majority of significant correlations occurs with genes of the carotenoid biosynthesis pathway in both vegetative and fruit tissues. These co-expression analyses confirm the importance of SlG2 and SlG3 in the tomato isoprenoid network and suggest that both isoforms are important to meet the demand of GGPP in chloroplasts and chromoplasts.

**Figure 3.**
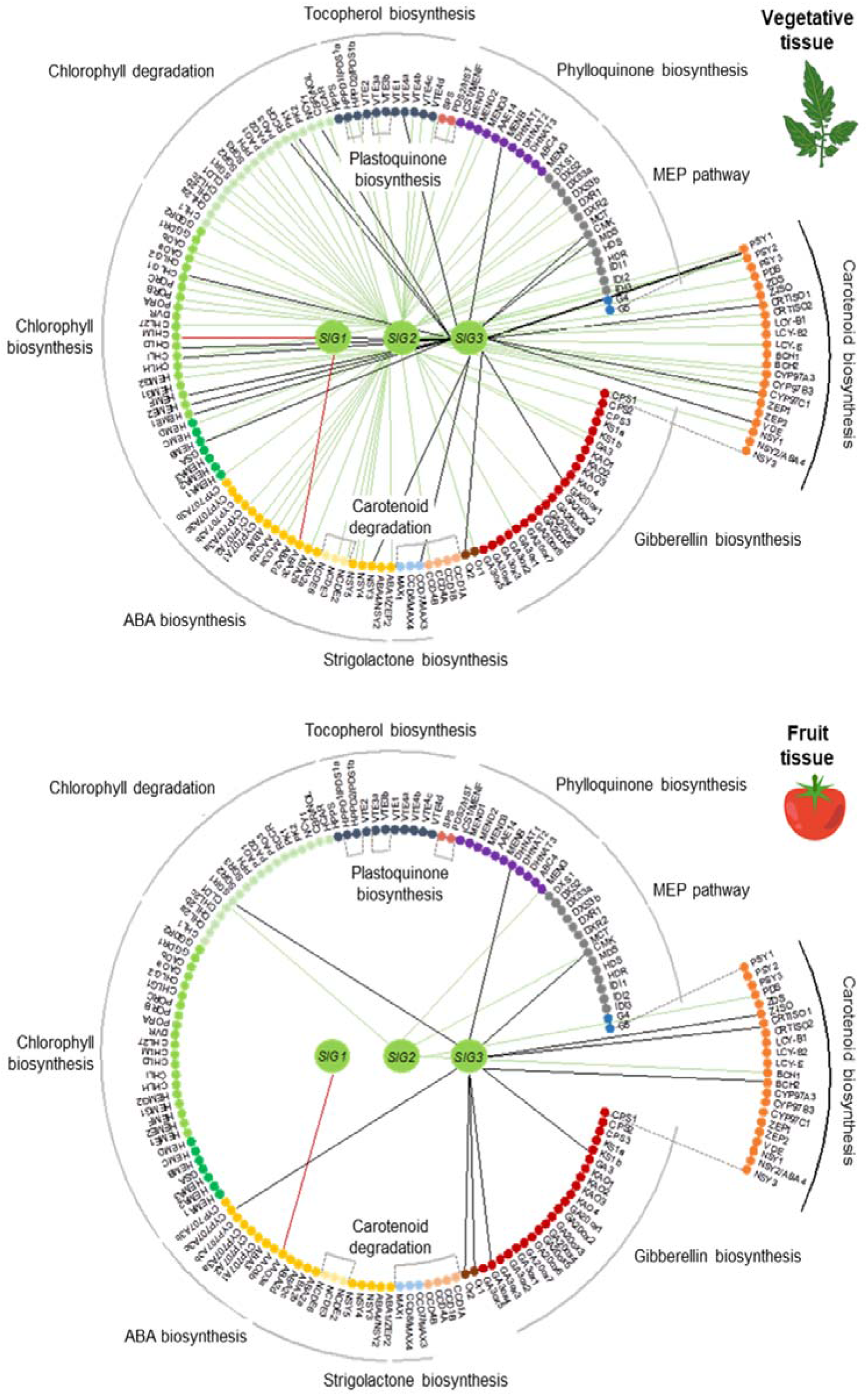
Gene co-expression analysis of tomato genes encoding plastidial GGPPS isoforms. Positive co-expression relationships (*ρ*≥0.55) are depicted in tissue-specific networks as edges. *SIG1, SIG2* and *SIG3* are depicted as central green nodes. Surrounding smaller nodes represent genes from the indicated isoprenoid pathways. Red, green and black edges indicate positive co-expression with *SIG1, SIG2* and *SIG3* genes, respectively. See Table S2 for gene accessions and Table S3 for *ρ* values.

In tomato, carotenoids contribute to mycorrhizal associations, photoprotection, and fruit pigmentation and, hence, the levels of these GGPP-derived metabolites increase during root mycorrhization, seedling de-etiolation, and fruit ripening. In agreement with the rate-determining role of PSY for carotenoid synthesis (Fraser et al., 2002), the expression levels of PSY-encoding genes also increase during such carotenoid-demanding developmental processes. By using real-time quantitative PCR (qPCR) analysis, we experimentally confirmed the up-regulation of *PSY1* during fruit ripening and *PSY3* in mycorrhized roots (Figure 4). Furthermore, we found that the *PSY2* gene was more strongly upregulated than *PSY1* during tomato seedling de-etiolation (Figure 4). Using the same samples, we observed that only *SlG1* was upregulated during root mycorrhization, similarly to the expression pattern observed for *PSY3* (Figure 4). During fruit ripening, *SlG2* and, to a lower extent, *SlG3* were up-regulated, but not as much as *PSY1* (Figure 5). *SlG2* was also the most strongly upregulated GGPPS-encoding gene during seedling de-etiolation, paralleling *PSY2* induction. Interestingly, *SlG3* and *PSY1* were also induced with a similar profile during this process, even though induction levels were much lower than those observed for *SlG2* and *PSY2* (Figure 4). Together, these data suggest that SlG1 might provide GGPP for PSY3 to produce carotenoids in roots, particularly when needed during mycorrhization, whereas both SlG2 and SlG3 would be required in leaves and fruits to support carotenoid production for photosynthesis (mostly via PSY2) and fruit pigmentation (via PSY1).

**Figure 4.**
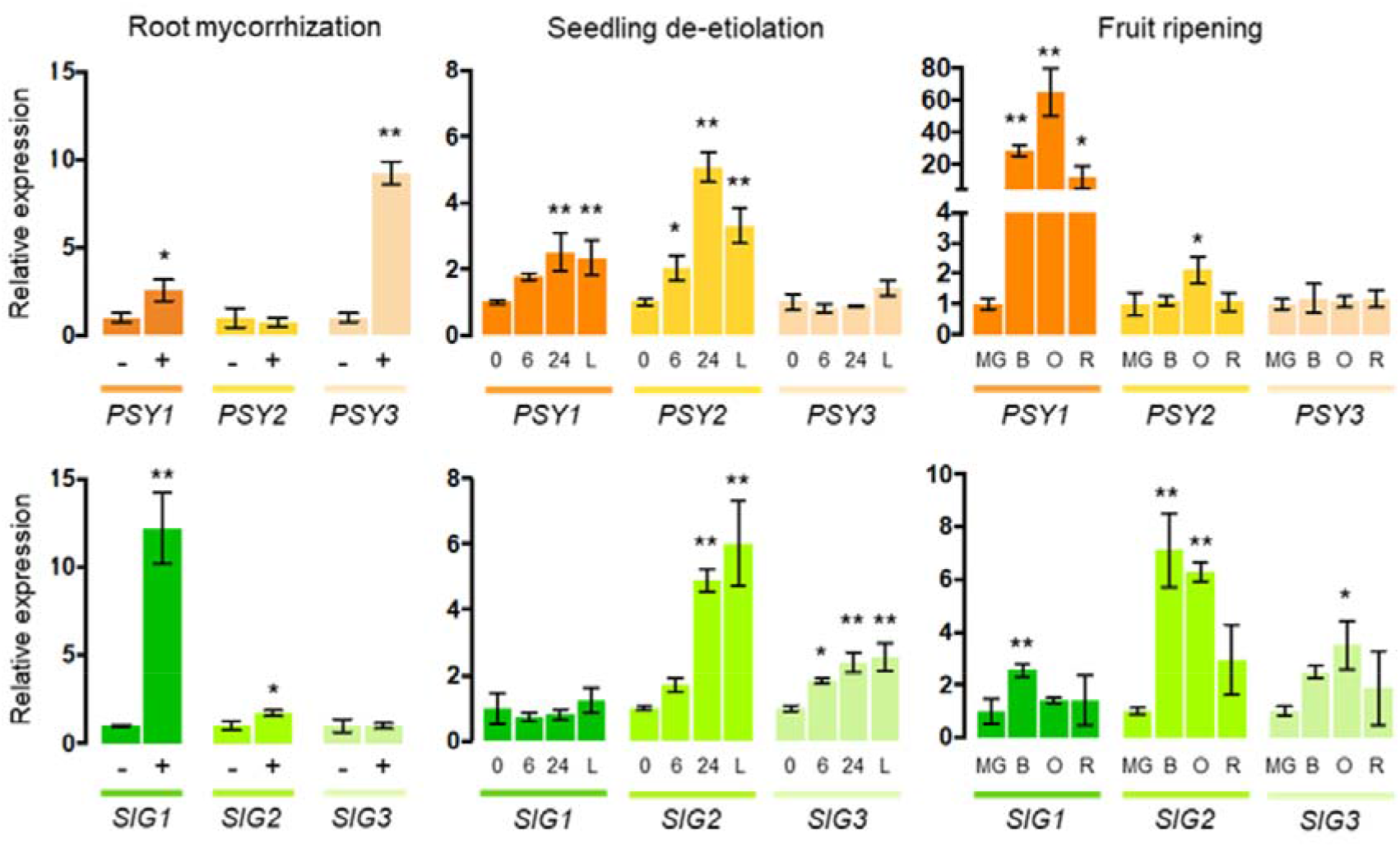
Expression profiles of genes encoding tomato PSY and GGPPS paralogs during processes involving increased carotenoid production. First column corresponds to non-mycorrhized (−) and mycorrhized roots (+) at 6 weeks post-inoculation. Transcript levels were normalized using the tomato *EXP* gene and are shown relative to untreated root samples. Central column samples correspond to 7-day-old dark-grown seedlings at 0, 6 and 24 h after exposure to light and to seedlings continuously grown in the light (L). Transcript levels were normalized to the *EXP* gene and are represented relative to etiolated (0 h) samples. Third column depicts different fruit ripening stages: MG, mature green; B, breaker; O, orange; and R, red ripe. Levels were normalized using *ACT4* and are shown relative to MG samples. Expression values represent the mean±SD of three independent biological replicates (n=3). Asterisks indicate statistically significant differences relative to untreated (−), etiolated (0 h) or MG samples (t-test or one-way ANOVA with Dunnett’s multiple comparisons test, *p<0.05, **p<0.01).

**Figure 5.**
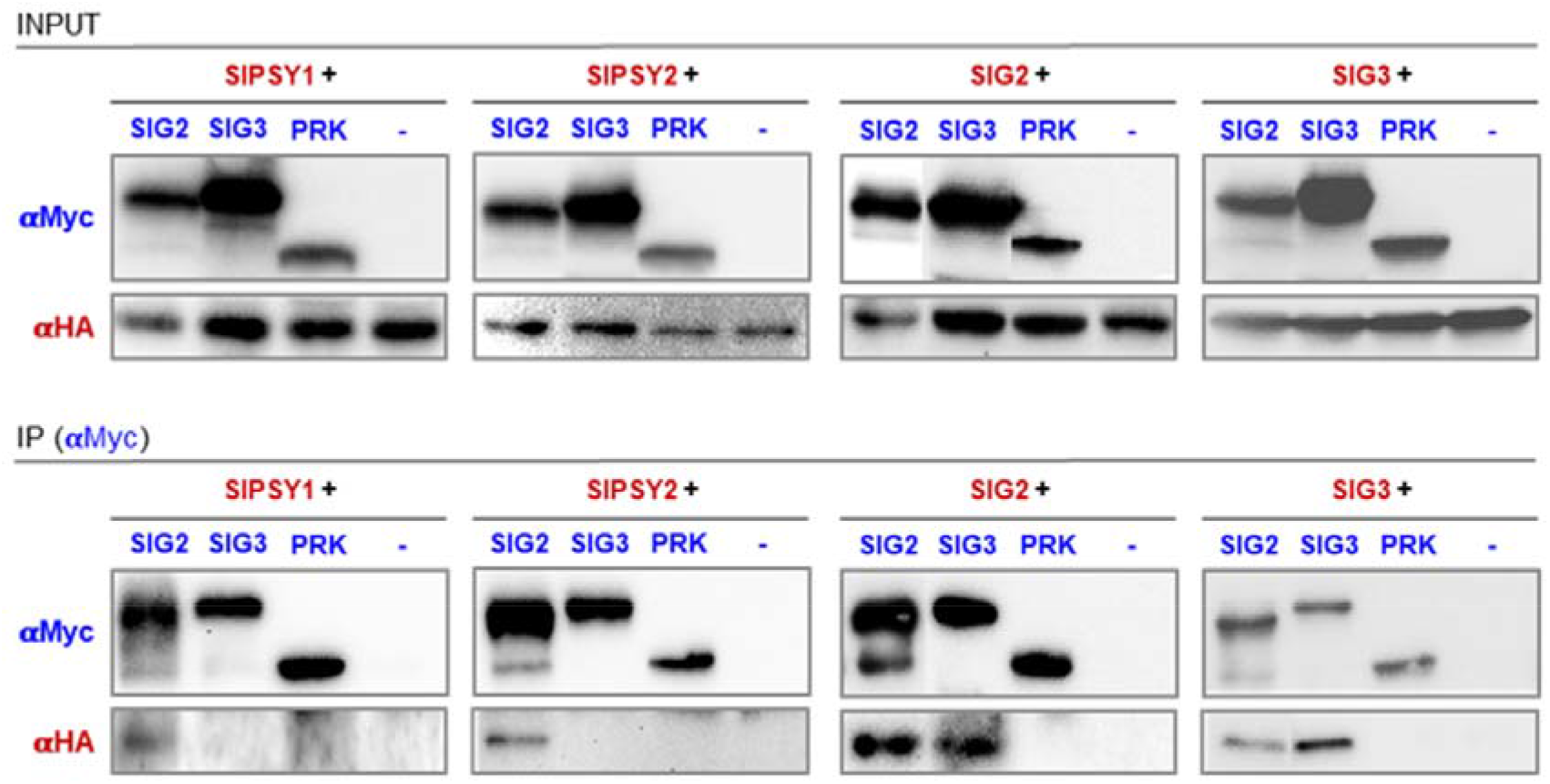
Co-immunoprecititation analyses. *N. benthamiana* leaves were co-agroinfiltrated with the indicated proteins tagged with C-terminal Myc (in blue) or HA (in red) epitopes. Controls agroinfiltrated only with the HA-tagged protein as indicated as (−). A fraction of the protein extracts (INPUT) was used to test protein production by immunoblot analyses using antibodies against Myc (αMyc) and HA (αHA). After immunoprecipitation (IP) of the remaining protein extracts using aMyc, samples were used for immunoblot analyses with αMyc (to confirm successful IP) and αHA (to detect the presence of co-immunoprecipitated HA-tagged proteins).

### SlG2, but not SlG3, can interact with PSY1 and PSY2

A coordinated role for SlG1 and PSY3 in mycorrhization has been already proposed (Stauder et al., 2018) but the possible connection between the other plastidial GGPPS and PSY isoforms remains unclear. GGPPS proteins can physically interact with PSY and other enzymes catalyzing both upstream and downstream biosynthetic steps in the plastids of different plant species (Dogbo and Camara, 1987; Camara, 1993; Maudinas et al., 1977a; Fraser et al., 2000; Ruiz-Sola et al., 2016a; Zhou et al., 2017; Camagna et al., 2019; Wang et al., 2018). This mechanism may facilitate channeling of precursors towards specific groups of plastidial isoprenoids. In particular, protein complexes containing both GGPPS and PSY enzymes were isolated from tomato chloroplasts and fruit chromoplasts (Maudinas et al., 1977a; Fraser et al., 2000). Nevertheless, the specific isoforms forming these protein complexes were never identified. Given the co-regulation of *SlG2* and *SlG3* with *PSY1* and *PSY2* genes in chloroplasts (i.e. photosynthetic tissues) and chromoplasts (i.e. fruits), we decided to test possible interactions of these isoforms (Figure 5). Constructs harbouring C-terminal Myc-tagged GGPPS and HA-tagged PSY sequences were combined and transiently co-expressed in *N. benthamiana* leaves. After 3 dpi, the recombinant SlG2-Myc and SlG3-Myc proteins were immunoprecipitated with αMyc antibody and the presence of PSY1-HA or PSY2-HA proteins in the samples was detected by immunoblot analyses using αHA antibody. As negative control, we used a Myc-tagged version of Arabidopsis phosphoribulokinase (PRK-Myc), a stromal enzyme of the Calvin cycle. Using this approach, we found that both PSY1-HA and PSY2-HA could be co-immunoprecipitated with SlG2-Myc, suggesting that they are present in the same complexes *in vivo* (Figure 5). By contrast, none of these PSY isoforms could be detected in the samples co-immunoprecipitated with either SlG3-Myc or PRK-Myc. The same Myc-tagged SlG2 and SlG3 proteins used in these experiments were able to co-immunoprecipitate their HA-tagged counterparts (Figure 5). This result, consistent with the ability of GGPPS proteins to form homodimers and also heterodimers, confirms that the observed lack of interaction of SlG3 with PSY enzymes was not due to SlG3-Myc having lost its capacity to interact with other proteins.

### Loss of function mutants defective in SlG3, but not those impaired in SlG2, show lower photosynthetic activity

To further understand the biological roles of SlG2 and SlG3, we generated CRISPR-Cas9 mutants defective in these enzymes (Figure 6). We designed two single guide RNAs (sgRNA) for each gene with the aim of creating deletions encompassing unique restriction sites for rapid screening (Figure 6A). Tomato plants of the variety MicroTom (MT) were transformed via *A. tumefaciens*, and two independent deletion alleles that created premature translation stop codons were selected for each gene and named *slg2-1*, *slg2-2*, *slg3-1* and *slg3-2* (Figure 6A) (Figures S6-S8). To confirm that the truncated proteins lacked GGPPS activity, we tested them in *E. coli* strains that synthesize the red carotenoid lycopene only when a source of GGPP is supplied (Figure 6B). While colonies harboring complete SlG2 or SlG3 enzymes produced lycopene, those transformed with the truncated versions of the tomato proteins were unable to synthesize GGPP for carotenoid biosynthesis (Figure 6B). Once confirmed that the selected mutant alleles produced non-functional proteins, homozygous lines without Cas9 were obtained and used for further experiments.

**Figure 6.**
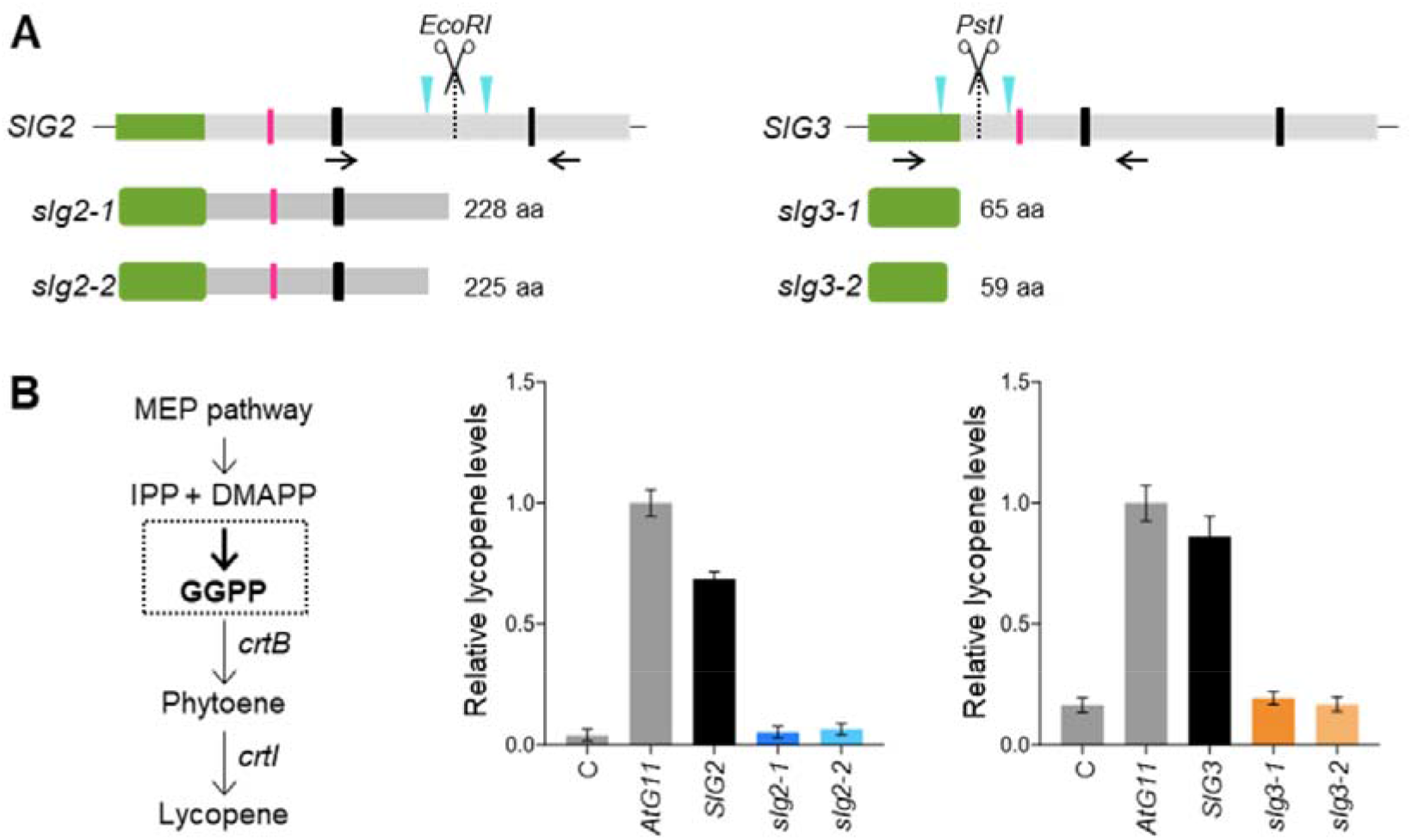
CRISPR-Cas9 mutagenesis of SIG2 and SIG3 genes. **(A)** Scheme representing the designed strategy to generate deletions on *SIG2* and *SIG3* genes and resulting proteins in selected mutant alleles (See Figures S6-S8 for further details). Green, pink and black boxes represent transit peptides, protein-protein interaction motifs, and catalytic domains (FARM and SARM), respectively. Blue arrowheads indicate the position of the designed sgRNAs encompassing specific restriction sites, and black arrows represent primer pairs used for genotyping. **(B)** Activity assays of WT and mutant GGPPS enzymes in *E. coli* strains expressing bacterial genes for lycopene biosynthesis (*crtB* and *crtl*) but lacking GGPPS activity. Lycopene production after transformation with constructs harboring the indicated sequences or empty plasmid controls (C) is shown relative to the levels obtained with the bona-fide GGPPS enzyme AtG11. Values represent the mean±SD of at least three independent transformants (n=3).

The most obvious phenotype among the selected lines was the pale color of *slg3* mutants compared to *slg2* alleles or azygous (WT) plants (Figure 7). This phenotype was clear in emerging and young leaves but it weakened as leaves grew and became mature (Figure 7A). The pale color correlated with significantly reduced levels of carotenoids and chlorophylls in young leaves of *sgl3-1* and *sgl3-2* lines compared to those of *sgl2* mutant alleles, which exhibited WT levels of photosynthetic pigments (Figure 7B) (Table S4). Tocopherols, other GGPP-derived plastidial isoprenoids, were also reduced in young leaves from SlG3-defective plants compared to WT or *slg2* samples (Table S4). This change was statistically significant in the *slg3-2* allele (Figure 7B) but also when samples from both *slg3-1* and *slg3-2* alleles were considered as a whole (Figure 8). By contrast, similar levels of carotenoids, chlorophylls and tocopherols were detected in mature leaves of WT, *slg2* and *slg3* plants (Figure 7B) (Table S4).

**Figure 7.**
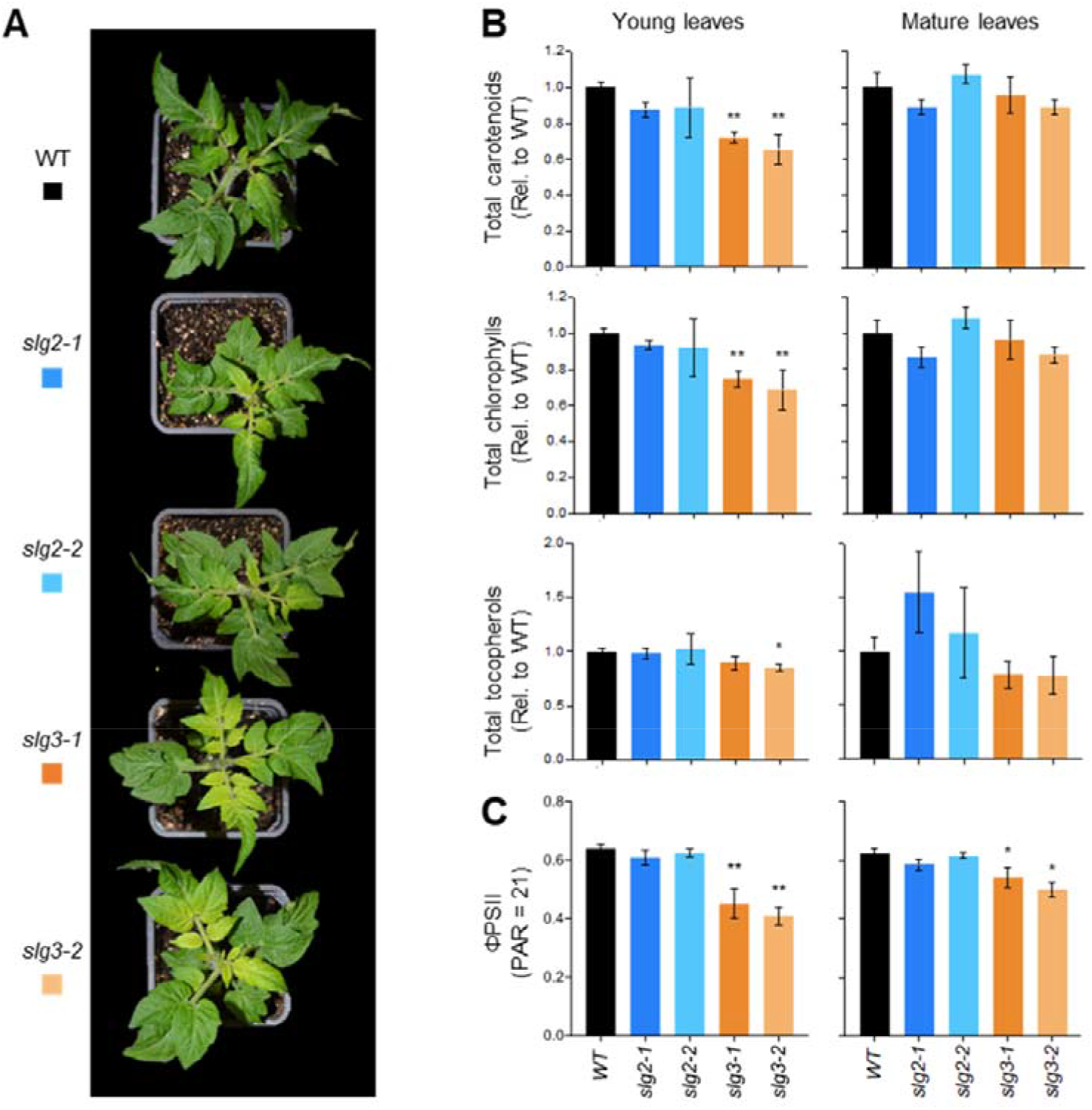
Leaf phenotypes of mutant lines defective in SIG2 or SIG3. **(A)** Representative images of 4-week-old plants of the indicated lines. **(B)** Relative levels of total carotenoids, chlorophylls and tocopherols in young and mature leaves of WT and mutant lines. Values are represented relative to WT levels and they correspond to the mean±SD of at least three independent biological replicates (n=3). See Table S4 for absolute values. **(C)** ϕPSII in young and mature leaves of the indicated lines. Values represent the mean±SD of three independent biological replicates (n=3). In all cases, asterisks indicate statistically significant differences among means relative to WT samples (one-way ANOVA with Dunnett’s multiple comparisons test, *p<0.05, **p<0.01).

**Figure 8.**
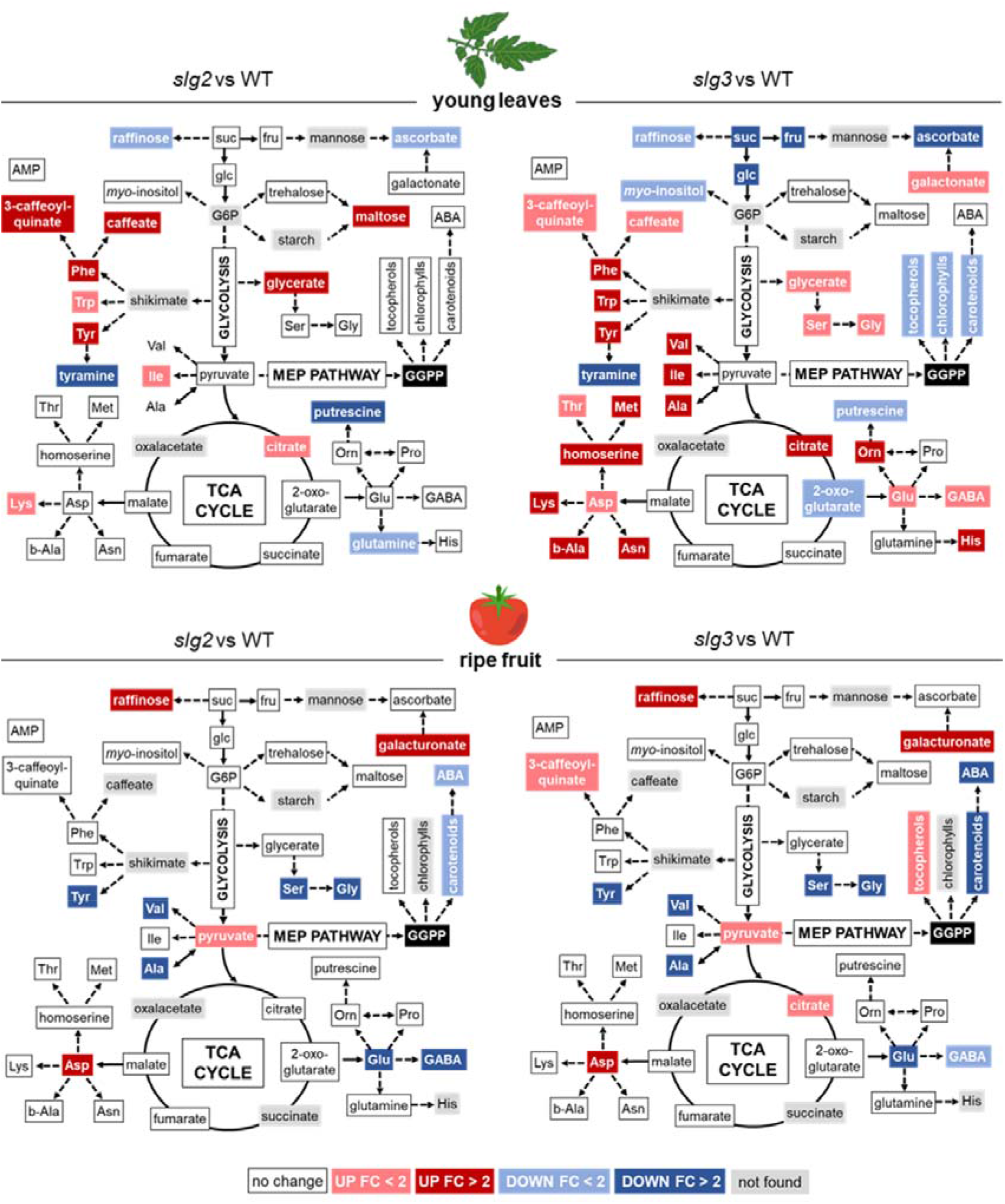
Metabolic changes in *slg2* and *slg3* mutants. Colors represent statistically significant fold-change (FC) values (t-test, p<0.05) of metabolite levels in young leaves or ripe fruit (B+10) from mutant plants relative to those in WT controls. See Table S5 (leaves) and S7 (fruit) for quantitative data.

To test whether the reduced accumulation of carotenoids and other photosynthesis-related isoprenoids in *slg3* lines had an impact on photosynthesis, we quantified effective quantum yield of photosystem II (ϕPSII) in both young and mature leaves from WT and mutant plants (Figure 7C). A 30% reduction in ϕPSII was observed in young leaves from *slg3* plants compared to those of WT or *slg2* lines, consistent with the *slg3-*specific reduction of GGPP-derived metabolites. Despite similar levels of photosynthetic pigments accumulated in the mature leaves of all genotypes tested, ϕPSII was slightly reduced in those from the *slg3* mutants relative to WT and *slg2* lines (Figure 7C). This result suggests that impaired production of photosynthesis-related isoprenoids in SlG3-defective young leaves might have an impact on photosynthesis that lasts even after recovering normal pigment levels in mature leaves.

We further explored possible effects that the loss of SlG2 or SlG3 function might have on other metabolic pathways using the same samples of young leaves used for isoprenoid and ϕPSII determination (Figure 8) (Tables S5 and S6). GC-MS metabolite profiling showed strongly decreased levels of sucrose, glucose and fructose in SlG3-defective leaves, likely due to photosynthetic impairment. Mutant *slg3* leaves also displayed increased levels of amino acids derived from glycerate (Ser and Gly), shikimate (Phe, Trp and Tyr), pyruvate (Val, Ile and Ala), 2-oxoglutarate (Glu, Orn, His and GABA) and malate (Asp, Asn, Lys, Thr, Met, homoserine and beta-alanine). In line with some of these amino acid changes, SlG3-defective leaves displayed altered accumulation of tricarboxylic acid cycle-related intermediates (citrate and 2-oxoglutarate). Only a few common changes were detected in both *slg2* and *slg3* leaves. They included a decrease in putrescine and ascorbate levels (more pronounced in *slg3* leaves), as well as an altered accumulation of metabolites produced by the plastidial shikimate pathway, including the above mentioned aromatic amino acids and phenylpropanoid derivatives such as caffeate and 3-caffeoyl-quinate (Figure 8) (Tables S5 and S6). The levels of the carotenoid-derived hormone ABA were similar in WT and mutant samples (Figure 8 and Table 2).

**Table 2.** ABA levels in GGPPS-defective leaves and fruit. Values (μg/g dry weight) correspond to the mean ± SD of four independent samples (n=4). Statistically significant changes in mutants compared to WT samples (t-test, p<0.01) are indicated in bold.

|  | Young leaves | B+10 fruit |
| --- | --- | --- |
| WT | 1.67 ± 0.19 | 0.63 ± 0.13 |
| *slg2-1* | 1.69 ± 0.10 | 0.55 ± 0.12 |
| *slg2-2* | 1.98 ± 0.39 | **0.30 ± 0.08** |
| *slg3-1* | 1.96 ± 0.09 | **0.16 ± 0.04** |
| *slg3-2* | 1.61 ± 0.29 | **0.08 ± 0.01** |

### Ripening-associated fruit pigmentation is altered in *slg2* and*slg3* mutants in correlation with their carotenoid profile

Lines with reduced levels of plastidial GGPPS activity also showed alterations in reproductive development (Figure 9). Flowering time was similar in WT, *slg2* and *slg3* plants (Figure 9A). However, pigmentation changes associated to fruit ripening were visually delayed in mutant fruits (Figure 9B). Tomato fruits reach their final size at the mature green (MG) stage and then start to ripe. The first visual symptoms of ripening define the breaker (B) stage, when chlorophyll degradation and carotenoid biosynthesis change the fruit color from green to yellow (Figure 9C). As ripening advances, accumulation of orange and red carotenoids (β-carotene and lycopene, respectively) progressively change the fruit color and define the orange (O) and eventually red (R) stages (Figure 9C).

**Figure 9.**
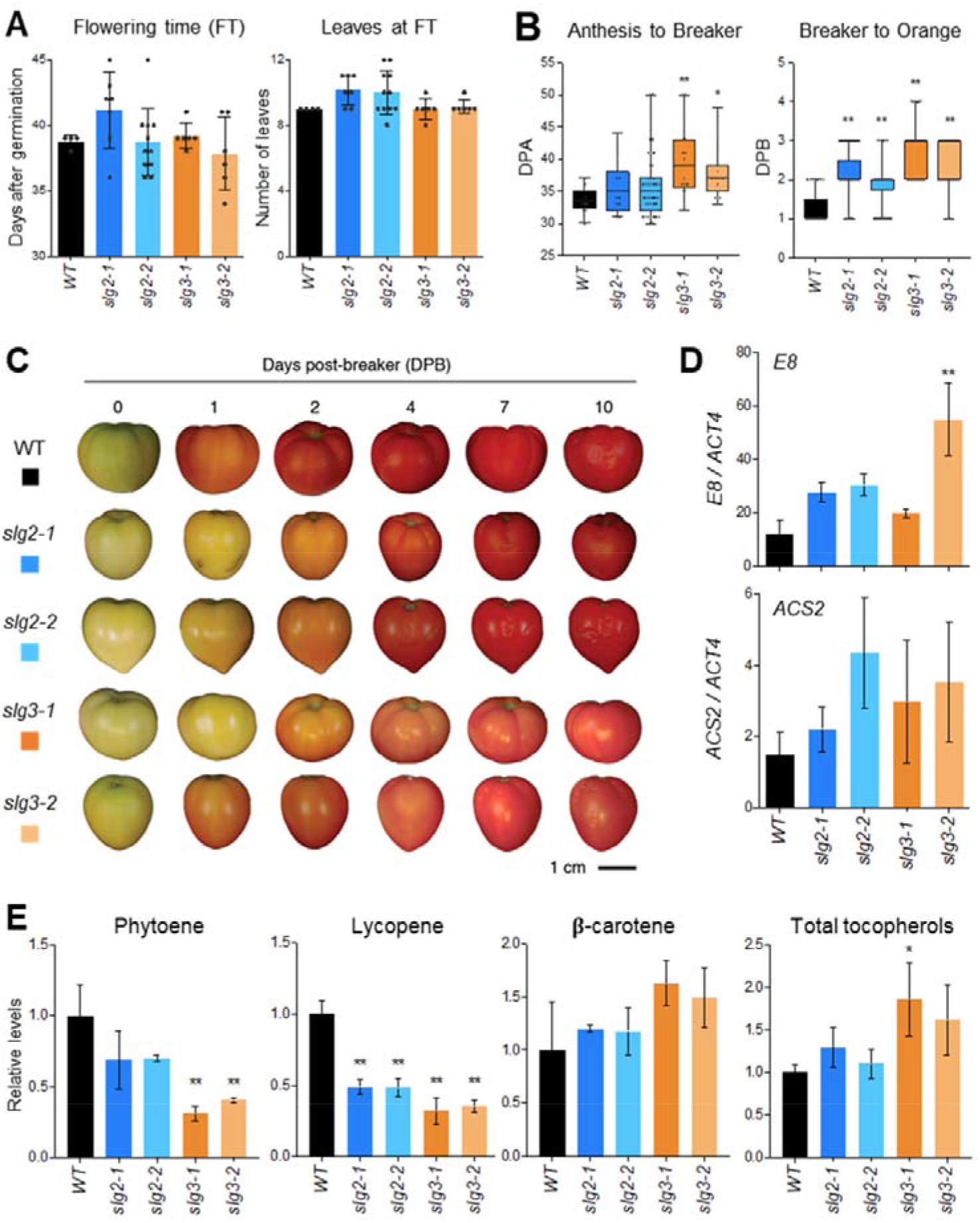
Flowering and fruit phenotypes of mutant lines defective in SIG2 or SIG3. **(A)** Flowering time measured as days after germination (left) or number of leaves (right). Values correspond to the meamSO of at least n=4 independent biological replicates. **(B)** Number of days to reach the indicated ripening stages represented as days post-anthesis on-vine (DPA, left) and days post-breaker off-vine (DPB, right). **(C)** Representative images of fruit from WT and mutant lines harvested at the breaker stage. **(D)** RT-qPCR analysis of *E8* and *ACS2* transcript levels in B+10 fruits of the indicated lines. Expression values were normalized using *ACT4* and represent the mean±SD of n=3 independent biological replicates. **(E)** Relative levels of individual carotenoids (phytoene, lycopene and β-carotene) and total tocopherols in B+10 fruits of WT and mutant lines. Values are represented relative to those in WT samples and correspond to the mean±SD of n=3 independent biological replicates. In all plots, asterisks indicate statistically significant differences among means relative to WT samples (one-way ANOVA with Dunnett’s multiple comparisons test, *p<0.05, **p<0.01).

The time from anthesis to B was similar in WT and SlG2-defective fruits but it was longer in the *slg3-1* and *slg3-2* mutants (Figure 9B) (Figure S9). Fruits from lines defective in SlG3, but also those defective in SlG2, showed a pigmentation delay in the transition from B to O. The delay was observed both on-vine (i.e. in fruits attached to the plant) and off-vine (i.e. in fruits detached from the plant at the B stage) (Figure 9B) (Figure S9). Both on-vine and off-vine measurements revealed that *slg2* mutants also took longer to reach the R stage compared to WT fruits (Figure S9), whereas *slg3* mutants did not reach a proper R stage as they developed a dark-orange color when ripe and never turned fully red (Figure 9C).

We next analyzed the levels of individual carotenoids and other GGPP-derived plastidial isoprenoids in fruits at MG (Figure S10) and B+10, i.e. 10 days after B (Figures 8 and 9E) (Table S4). WT and mutant fruits showed similar levels of carotenoids, chlorophylls and tocopherols at the MG stage, but clear differences were detected in ripe (B+10) fruits. Phytoene and lycopene were significantly decreased in all mutants, although the impact was higher in the case of *slg3* fruits. No significant differences were found for β-carotene, although the levels of this orange carotenoid tended to be higher in *slg3* mutants. This, together with the lower levels of the red carotenoid lycopene may explain the dark orange color of B+10 *slg3* fruits (Figure 9C). Tocopherols also showed a trend towards higher abundance in SlG3-deficient fruits, a change that was statistically significant in the *slg3-1* allele (Figure 9E) or when *slg3-1* and *slg3-2* samples were considered together (Figure 8).

Unlike that observed in young leaves, ABA levels were reduced in B+10 fruits of *slg2* and, most strongly, *slg3* mutants compared to WT controls (Figure 8 and Table 2). A role for ABA in promoting tomato fruit ripening has been proposed based on the analysis of mutants or external application of hormones and inhibitors (Seymour et al., 2013; Quinet et al., 2019). While decreased ABA levels in *slg2* and *slg3* fruits might be expected to result in slower ripening, analysis of ripening marker genes such as *E8* and *ACS2* (Figure S4) (Estornell et al., 2009; Llorente et al., 2016; D’Andrea et al., 2018) did not show a substantial ripening delay in mutant fruits (Figure 9D). At the level of primary metabolites, B+10 fruits from both *slg2* and *slg3* mutants exhibited increased levels of raffinose, galacturonate, pyruvate and Asp, and lower levels of Ser, Gly, Tyr, Val, Ala, Glu and GABA than WT controls (Figure 8) (Tables S7 and S8). The observation that metabolite changes are very similar in both *slg2* and *slg3* fruits suggests that SlG2 and SlG3 have redundant roles during fruit ripening. The strongest decrease in carotenoids and derived ABA detected in the *slg3* mutant further indicates that SlG3 likely has a more prominent role for the production of ripening-related carotenoids.

### Double mutants defective in both SlG2 and SlG3 are not viable

To assess the impact of simultaneous disruption of both *SlG2* and *SlG3* genes, alleles *slg2-2* and *slg3-1* were crossed using the former as female parent and the latter as male parent or vice versa. Double heterozygous F1 plants from each cross were allowed to self-pollinate and the resulting seeds were used to screen the F2 population for double homozygous plants, which were expected to occur at a frequency of 6.25% (1 in 16). PCR genotyping of 200 F2 seedlings (100 from each cross), however, found no double homozygous mutants (Table 3). Interestingly, single azygous plants (i.e. *SlG2 slg2 slg3 slg3* and *slg2 slg2 SlG3 slg3*) were found at lower frequencies than expected (Table 3), suggesting a gene dosage effect on the production of essential GGPP-derived isoprenoids. Hence, the absence of both *SlG2* and *SlG3* appears to result in a lethal phenotype that is partially rescued by incorporating one copy of any of these two genes (as in single azygous plants) and fully rescued when two copies are present in the genome (as in double heterozygous or single homozygous mutants). These results, together with the similar expression levels of both genes in developing tomato seeds (Figure S4), suggest that SlG2 and SlG3 contribute similarly and additively to embryo or/and seed development.

**Table 3.** Expected and observed frequencies of the F2 population from the crosses of *slg2-2* and *slg3-1*. Mutant alleles are marked in red. Percentages in bold indicate observed frequency lower than expected.

| Genotypes | Expected | Observed | Plants |
| --- | --- | --- | --- |
| G2g2 G3g3 | 25% | 26% | 52 |
| G2g2 G3G3 | 12.5% | 13% | 26 |
| G2G2 G3g3 | 12.5% | 17.5% | 35 |
| **g2g2** G3g3 | **12.5%** | **9%** | **18** |
| G2g2 **g3g3** | **12.5%** | **8%** | **16** |
| **g2g2** **g3g3** | **6.25%** | **0%** | **0** |
| **g2g2** G3G3 | 6.25% | 8.5% | 17 |
| G2G2 **g3g3** | 6.25% | 7% | 14 |
| G2G2 G3G3 | 6.25% | 11% | 22 |

## DISCUSSION

GGPP synthesis and allocation to different downstream pathways in plants is controlled by GGPPS enzyme families, whose members are regulated at transcriptional and post-transcriptional level (Beck et al., 2013; Ruiz-Sola et al., 2016a, 2016b; Zhou et al., 2017; Stauder et al., 2018; Wang et al., 2018). Several efforts have been done since the late 90’s to characterize the GGPPS protein family in Arabidopsis, establishing the fundamental basis for our knowledge of the regulation of GGPP biosynthesis in plants (Zhu et al., 1997a, 1997b; Okada et al., 2000; Beck et al., 2013; Nagel et al., 2015; Ruiz-Sola et al., 2016a, 2016b; Wang et al., 2016b). In this model plant, two GGPPS paralogs (AtG2 and AtG11) have been shown to produce GGPP and to be targeted to plastids. However, only AtG11 appears to be required for the production of plastidial isoprenoids (Beck et al., 2013; Nagel et al., 2015; Ruiz-Sola et al., 2016a, 2016b). The gene encoding AtG11 is ubiquitously expressed at high levels and can generate long transcripts encoding the plastid-targeted isoform but also short transcripts encoding a cytosolic enzyme that retains enzymatic activity and is essential for embryo development (Ruiz-Sola et al., 2016b). The production of GGPP has also been studied in a few crop plants (Wang and Dixon, 2009; Zhang et al., 2015; Zhou et al., 2017; Wang et al., 2018, 2019). Similar to Arabidopsis, rice and pepper contain only one enzymatically active GGPPS isoform localized in plastids, named OsGGPPS1 (OsG1 in short) and CaGGPPS1 (CaG1), respectively (Zhou et al., 2017; Wang et al., 2018). Strikingly, little information is available on the tomato GGPPS family despite this species being a well-established model plant that accumulates high amounts of GGPP-derived metabolites of human interest such as carotenoids in fruits. Here we have gathered information on the tomato GGPPS family, providing evidence of a coordinated role of two plastidial isoforms (SlG2 and SlG3) on supplying GGPP to produce carotenoids and other isoprenoids essential for photosynthesis, fruit pigmentation, and seed viability.

All the GGPPS paralogs identified here (SlG1-5) harbored the seven domains known to be highly conserved in SC-PTs (Koike-Takeshita et al., 1995) and all other features known to be required for GGPPS activity (Figure S1). Moreover, they were differentially located in plastids, cytosol and mitochondria (Figure 2A), consistent with the need of GGPPS isozymes in these cell compartments to feed the different organelle-specific isoprenoid pathways (Bick and Lange, 2003; Beck et al., 2013; Ruiz-Sola et al., 2016a, 2016b; Zhou et al., 2017; Wang et al., 2018). We confirmed that the three plastid-targeted GGPPS homologs (SlG1-3) produce GGPP with similar kinetic parameters (Figure 2B and Table 1) and an optimal pH around 7.5 (Figure S3), similar to the stromal pH (Höhner et al., 2016). SlG5-GFP was targeted to mitochondria, whereas the GFP-fused SlG4 protein exhibited a major preference for cytosolic location but it was also observed in the chloroplasts of some *N. benthamiana* cells (Figure 2A) (Figure S2). Further illustrating a complex localization pattern for the SlG4 protein, fusion of its N-terminal sequence to GFP generated a mitochondrial fluorescence pattern when expressed in Arabidopsis protoplasts (Zhou and Pichersky, 2020). Multiple subcellular localization of many isoprenoid biosynthetic enzymes has been widely described (Cunillera et al., 1997; Phillips et al., 2008; Sapir-Mir et al., 2008; Ruiz-Sola et al., 2016b). In many cases, including Arabidopsis *AtG11* (Ruiz-Sola et al., 2016b), the production of two differentially-targeted proteins from the same gene is due to a weak transcription initiation site on the corresponding gene, giving rise to some transcripts that lack the first ATG codon of the coding sequence. Thus, translation starts from a second in-frame ATG codon producing a shorter protein lacking the N-terminal region (which usually corresponds to the organelle-specific transit peptide). A similar transcriptional regulation could generate long versions of the SlG4 protein that would be targeted to chloroplasts or mitochondria, as well as short versions that would be retained in the cytosol. Despite the clear plastidial localization observed here (Figure 2A) and elsewhere (Zhou and Pichersky, 2020) for GFP fusions of the SlG1-3 isoforms, we cannot exclude that shorter versions of these proteins could also be produced *in vivo*, paralleling that observed in the case of AtG11 (Ruiz-Sola et al., 2016b). This possibility is strongly supported by the observation that no double mutants defective in both SlG2 and SlG3 could be identified (Table 3), suggesting that the loss of these two GGPPS isoforms might cause an embryo-lethal phenotype similar to that reported in Arabidopsis mutants lacking the short (i.e. cytosolic) version of AtG11 (Ruiz-Sola et al., 2016b). Indeed, several M residues can be found in the N-terminal region of both SlG2 and SlG3 enzymes (Figure S8); they could be used as alternative translation start sites to produce catalytically active GGPPS enzymes with an absent or shorter (i.e. dysfunctional) plastid-targeting domain.

SlG1-3 are the only GGPPS-related sequences in tomato shown to actually produce GGPP as their main product (Figure 2B) (Zhou and Pichersky, 2020). The presence of these three GGPPS isoforms in tomato plastids may be caused by the acquisition of specific roles during evolution (subfunctionalization). Besides localization in distinct subcellular compartments, new functions can be acquired either through a differential spatio-temporal gene expression, triggered by developmental and environmental signals, or through specific interactions with other proteins. Mining of public tomato gene expression databases, GCN analyses and qPCR assays led us to conclude that SlG1 is likely contributing to carotenoid biosynthesis in roots together with PSY3. This conclusion is supported by a recent study showing that the expression of *PSY3* and *SlG1* coordinately responds to tomato root mycorrhization and phosphate starvation (Stauder et al., 2018). The SlG1-PSY3 tandem might be channeling the flux of MEP-derived precursors towards the synthesis of carotenoid-derived molecules that are crucial for the establishment of symbiosis, such as strigolactones and apocarotenoids, including C13 α-ionol and C14 mycorradicin (Stauder et al., 2018). The observation that *SlG1* was significantly co-expressed with genes from the ABA synthesis pathway in vegetative and fruit tissues (Figure 3) is consistent with a possible involvement of this GGPPS isoform in the production of ABA under certain environmental conditions. *SlG1* expression was also found to be induced in tomato leaves after herbivore-feeding or treatment with jasmonates, correlating with an increase in the emissions of GGPP-derived volatile isoprenoids (Ament et al., 2006). It is therefore possible that this isoform specializes in the formation of GGPP in response to biotic interactions not only in roots, but also in other plant tissues such as leaves.

Unlike *SlG1*, *SlG2* and *SlG3* are constitutively expressed, with *SlG3* being the paralog with the highest expression level in all plant tissues (Figure S4). In green tissues, *SlG2* is more strongly co-expressed than *SlG3* with genes from photosynthesis-related isoprenoid pathways, exhibiting higher correlations than with genes involved in carotenoid biosynthesis (Figure 3). Additionally, *SlG2* was much more upregulated than *SlG3* during seedling de-etiolation (Figure 4) and leaf development (Figure S4), when an enhanced production of carotenoids and other photosynthesis-related isoprenoids contributes to assemble a functional photosynthetic machinery. *SlG2* was also much more induced that *SlG3* during fruit ripening, when carotenoid biosynthesis is boosted. The production of carotenoids during ripening relies on the PSY1 isoform, whose strong induction during fruit ripening has been recently explained as a probable consequence of its poor enzyme activity (Cao et al., 2019). While we did not find a significant correlation of *SlG2* or *SlG3* with *PSY1* expression in fruits, *PSY1* and *SlG2*, but not *SlG3*, are coordinately regulated by FUL and RIN transcription factors that control the expression of ripening-related genes, including many of the MEP and carotenoid pathways (Fujisawa et al., 2013, 2014). All these expression data show that *SlG2* expression is more responsive to increased demands of precursors to support carotenoid biosynthesis. In contrast, *SlG3* expression is higher and does not change so much, suggesting a house-keeping role to maintain a continuous supply of GGPP in plastids for carotenoids and other isoprenoids. According to this model, SlG1 and SlG2 would help SlG3 to supply GGPP when needed. The very low and restricted expression level of *SlG1*, however, strongly suggests that SlG2 is the main helper isoform for SlG3 in chloroplasts and chromoplasts.

Besides diverging gene expression profiles, subfunctionalization of GGPPS paralogs might also involve isoform-specific interactions with other proteins. The interaction of different plant GGPPS enzymes with other proteins involved in isoprenoid biosynthesis has been demonstrated through different protein-protein interaction approaches (Dogbo and Camara, 1987; Camara, 1993; Maudinas et al., 1977b; Fraser et al., 2000; Ruiz-Sola et al., 2016a; Zhou et al., 2017; Camagna et al., 2019; Wang et al., 2018). The enzymatic properties of GGPPS proteins change to produce GPP upon heterodimerization with members of the GPP synthase small subunit type I (SSU-I) subfamily (Orlova et al., 2009; Wang and Dixon, 2009). Thus, the tomato SSU-I protein (Solyc07g064660) interacts with SlG1-3 enzymes in plastids to change their product specificity from GGPP to predominantly GPP (Zhou and Pichersky, 2020). Other protein-protein interactions forming multienzymatic complexes appear to be particularly important for metabolic channeling of GGPP into the different pathways that consume it, as its high amphipathicity makes it unlikely that GGPP could diffuse in monodisperse form (Camagna et al., 2019). In particular, spatial proximity of GGPPS and PSY enzymes appears to be a precondition for efficient phytoene synthesis, because PSY cannot access freely diffusible GGPP or time-displaced GGPP supply by GGPPS (Camagna et al., 2019). Arabidopsis AtG11 and pepper CaG1 can directly interact with PSY proteins (Ruiz-Sola et al., 2016a; Camagna et al., 2019; Wang et al., 2018). We found that tomato SlG2, but not SlG3, is able to interact with PSY1 and PSY2 *in planta* (Figure 5). However, tomato SlG3 might deliver GGPP to PSY enzymes via heterodimerization with PSY-interacting SlG2 (Figure 5). An alternative possibility involves interaction with members of another catalytically-inactive SSU subfamily named type II (SSU-II). Similar to AtG11 and CaG1, OsG1 is the only GGPPS enzyme producing GGPP for carotenoid biosynthesis in rice. Strikingly, OsG1 does not interact with PSY but heterodimerizes with a SSU-II homolog named GGPPS Recruiting Protein, OsGRP (Zhou et al., 2017). While binding of SSU-I proteins changes the product specificity of GGPPS enzymes from GGPP to GPP (Orlova et al., 2009; Wang and Dixon, 2009; Zhou and Pichersky, 2020), interaction with the SSU-II protein OsGRP enhanced OsG1 catalytic efficiency to produce GGPP. Additionally, OsGRP binding delivered the OsG1 protein to a large protein complex in thylakoid membranes with enzymes involved in chlorophyll biosynthesis (Zhou et al., 2017). A plastid-targeted and catalytically inactive SSU-II protein was also shown to interact with pepper CaG1 and enhance its GGPP-producing activity (Wang et al., 2018). Interestingly, the pepper SSU-II protein also interacts with PSY, suggesting that it might stimulate both GGPPS activity and interaction with PSY (Wang et al., 2018). The tomato SSU-II homolog (Solyc09g008920) is an ubiquitously expressed and catalytically inactive protein targeted to plastids, as its rice and pepper counterparts (Zhou and Pichersky, 2020). In agreement with the conclusion that tomato, pepper and rice SSU-II homologs share similar functions, heterodimers formed between tomato SSU II and SlG1-3 catalyzed the formation of GGPP with improved efficiency (Zhou and Pichersky, 2020). It is possible that SlG3 might be delivered to PSY-containing protein complexes through interaction with tomato SSU-II. This protein might also enhance interaction of SlG2 with PSY isoforms, similar to that proposed for PSY-interacting CaG1 (Wang et al., 2018). A role for SSU-II in promoting interaction between GGPPS and PSY enzymes might be key to enhance carotenoid biosynthesis in plants.

The biochemical characterization of purified SlG2 and SlG3 enzymes together with the phenotypes of *slg2* and *slg3* mutants supported the conclusion that these are very similar enzymes with functionally interchangeable functions whose differential role as helper or house-keeping is basically determined by their expression pattern. Analysis of single tomato mutants defective in either SlG2 or SlG3 suggested that absence of any of the two individual enzymes decreases GGPP production and hence delays the production of photosynthetic isoprenoids in leaves (Figure 7) and carotenoids in fruits (Figure 9). Consistent with the higher expression levels, these phenotypes were stronger in *slg3* alleles. However, the effects of reduced isoprenoid synthesis could also be directly or indirectly detected in *slg2* plants. Thus, a carotenoid-associated pigmentation delay was more subtle but still detected in ripening fruit from *sgl2* alleles (Figure 9). Our HPLC analysis could not detect reduced levels of carotenoids or any other GGPP-derived isoprenoid in SlG2-defective leaves but the GC-MS analysis showed higher levels of all aromatic amino acids derived from the shikimate pathway (Trp, Tyr and Phe) as well as Phe-derived phenylpropanoids caffeate (caffeic acid) and 3-caffeoyl-quinate (chlorogenic acid) in both *slg2* and *slg3* mutant lines (Figure 8). This might be a physiological response to cope with photooxidative stress caused by lower levels of carotenoids in the mutants, as phenylpropanoids (including Phe-derived flavonoids and anthocyanins) can also function as photoprotective metabolites (Muñoz and Munné-Bosch, 2018). Reduced levels of well-known metabolites associated with oxidative stress such as ascorbate and putrescine in leaves from both mutant lines would also support this view. When levels of GGPP-derived photoprotective isoprenoids such as carotenoids and tocopherols fell below a certain threshold (and hence the decrease became detectable, as in young leaves of the SlG3-defective alleles), photosynthesis was eventually impacted (Figure 7C), causing sugar starvation and the subsequent metabolic changes observed only in the *slg3* mutant (Figure 8) (Tables S5 and S6). In agreement, the increased accumulation of most amino acids in *slg3* leaves suggested a high proteolytic activity to generate an alternative respiratory source, a likely response to sugar starvation derived from reduced photosynthesis and/or photooxidative stress (Araújo et al., 2011; Obata and Fernie, 2012; Galili et al., 2016). In ripe (i.e. non-photosynthetic) fruits, increased levels of 3-caffeoyl-quinate, citrate and tocopherols were detected only in *slg3* samples. The rest of metabolic changes detected in fruits were also similar in *slg2* and *slg3* lines (Figure 8) (Tables S7 and S8), again supporting the conclusion that these enzymes are redundant and interchangeable.

Because ABA is synthesized from carotenoids, its reduced levels in GGPPS-defective ripe fruits but not in leaves (Table 2) may be the result of a more substantial reduction in carotenoid contents in mutant fruit (Figure 9) compared to leaves (Figure 7) (Table S4). When the metabolic profile of ABA-defective B+10 fruit from *slg2* and *slg3* lines was compared with that of ripe fruit from other tomato lines with altered ABA levels, no correlation was found. For example, Glu and GABA were downregulated in our samples (Figure 8) (Tables S7 and S8) and in transgenic lines in which a constitutively silenced □-carotene desaturase (ZDS) prevents transformation of phytoene into lycopene and hence results in reduced ABA levels (McQuinn et al., 2020), but also in ABA-overaccumulating transgenic fruits with a constitutively expressed lycopene β-cyclase (LCYB) that converts lycopene into β-carotene (Diretto et al., 2020). Asp increased in ripe fruits with reduced ABA levels either by decreasing plastidial GGPPS (Figure 8) or ZDS activity (McQuinn et al., 2020) while it decreased in those with higher ABA levels (Diretto et al., 2020). By contrast, levels of Ser, Gly, Tyr, Val and Ala were decreased in ABA-defective *slg2* and *slg3* mutants (Figure 8) and ABA-accumulating LCYB overexpressors (Diretto et al., 2020) but increased In ZDS-silenced lines with lower ABA levels (McQuinn et al., 2020). Together, the results indicate that most of the observed amino acid changes in these mutant and transgenic fruits cannot be explained by altered ABA levels. Although reduced ABA might delay ripening (Seymour et al., 2013; Quinet et al., 2019), a slower production of carotenoids in GGPP-depleted mutant fruit could explain by itself the longer time required for these fruits to advance in color-based ripening phases (Figure 9B). Additionally, metabolic roles of SlG2 and SlG3 besides their GGPPS activity in plastids might play a role in fruits but also in developing seeds, hence explaining why we could not isolate a double *slg2 slg3* mutant (Table 3). If SlG2 and SlG3 play the same role in tomato that AtG11 plays in Arabidopsis, as it appears to be the case based on the data discussed above, it would be expected that the tomato double mutant was not rescued because an embryo-lethal phenotype results from an impaired activity of these enzymes in the cytosol (Ruiz-Sola et al., 2016b). Further experimental evidence would be required to confirm whether the use of alternative ATG codons to produce shorter (i.e. cytosolic) versions of otherwise plastidial GGPPS enzymes is a general mechanism in plants, and also to identify what is the product essential for embryo development supplied by the cytosolic enzymes (either alone or in combination with interacting proteins). In any case, the observation that the lethal phenotype is dose-dependent in an isoform-independent fashion (i.e. can be rescued by a single genomic copy of either *SlG2* or *SlG3*) reinforces our conclusion that SlG2 and SlG3 are enzymes with functionally interchangeable roles.

In summary, this work demonstrates that the bulk of GGPP production in tomato leaf chloroplasts and fruit chromoplasts relies on two redundant but cooperating GGPPS paralogs, SlG2 and SlG3. Additionally, the SlG1 isoform might contribute to GGPP synthesis in root plastids. This scenario contrasts with that described to date in other plant species such as *Arabidopsis thaliana*, rice or pepper, which produce their essential plastidial isoprenoids using a single GGPPS isoform. Deciphering how different plants regulate plastidial GGPP production and channeling will be useful for future metabolic engineering approaches targeted to manipulate the accumulation of specific groups of GGPP-derived isoprenoids without negatively impacting the levels of others. In the case of tomato carotenoids, understanding how GGPPS and PSY isoform pairs interact should allow to specifically improve the nutritional quality of fruits without interfering with other vital processes such as photoprotection, mycorrhization, or stress tolerance.

## MATERIALS AND METHODS

### Plant material and growth conditions

Tomato (*Solanum lycopersicum* var. MicroTom) plants were used for most experiments. Seeds were surface-sterilized by a 15 min incubation in 25 mL of 40% bleach containing a drop of Tween-20 followed by 3 consecutive 10 min washes with sterilized milli-Q water. Sterile seeds were germinated on plates with solid 0.5x Murashige and Skoog medium without vitamins or sucrose. The medium was supplemented with kanamycin (100 μg/mL) when required to select transgenic plants. After stratification at 4 °C in the dark for at least 3 days, plates were incubated in a climate-controlled growth chamber at 24 °C with a photoperiod of 10 h of darkness for 14 h of fluorescent white light at a photosynthetic photon flux density of 140 μmol m^−2^ s^−1^. After 1 to 2 weeks, old seedlings were then transferred to soil and grown under standard greenhouse conditions (14 h light at 27 ± 1 °C and 10 h dark at 22 ± 1 °C). For the analysis of flowering time, at least five independent plants of each genotype were used. Flowering time was assessed by counting the number of days from germination until the first flower was fully opened (anthesis) or the number of leaves in the plant at this first anthesis day.

For deetiolation experiments, seeds were sown on sterile water-soaked cotton in plastic containers. After stratification, seeds were exposed to fluorescent white light for 2-4 hours at 22°C to induce germination. The containers were then covered with a double layer of aluminum foil and kept in darkness at 22 °C. After one week, seedlings were exposed to light and samples were harvested after 0, 6 and 24 h. Control samples were germinated and grown under continuous light and collected at the 0 h time point. Leaf samples were collected from four-week-old plants. Young leaf samples correspond to growing leaflets from the fifth and sixth true leaves, and mature leaf samples correspond to fully expanded leaflets from the third or fourth leaf. Tomato fruit pericarp samples were collected at four ripening stages based on days post-anthesis (DPA) or days post-breaker (DPB): mature green (~30 DPA), breaker (~35 DPA), orange (~38-40 DPA) and red (~45-50 DPA or 10 DPB). Full seedlings, leaflets, and pericarp samples were frozen in liquid nitrogen immediately after collection, freeze-dried and stored at −80 °C. *Nicotiana benthamiana* plants were grown and used for transient expression assays (agroinfiltration) as previously described (Llorente et al., 2019).

### Constructs

Full-length cDNAs encoding SlG1-5 and PSY1-2 proteins without their stop codons were amplified by PCR and cloned via BP clonase into pDONR207 entry plasmid using Gateway (GW) technology (Invitrogen). Full-length sequences were then subcloned through an LR reaction into pGWB405 plasmid for subcellular localization assays, or into pGWB414 and pGWB420 plasmids for co-immunoprecipitation experiments. Constructs in pGWB405, pGWB414 and pGWB420 vectors harbor GFP, 3x-HA and 10x-Myc tags, respectively. These tag sequences are fused to the C-terminus of each cloned element and the expression module is controlled by the CaMV 35S promoter. For recombinant protein production in *E. coli*, SlG1-3 versions lacking the predicted transit peptide for plastid import were amplified from pGWB405 constructs, cloned into pDONR207 plasmid, and then subcloned into pET32-GW plasmid (fusing a 6x-His tag at the N-terminal end of the cloned fragments) under the control of the T7 promoter. pET32-GW constructs containing Arabidopsis AtG11 versions are described in (Ruiz-Sola et al., 2016b).

For CRISPR-Cas9-mediated disruption of *SlG2* and *SlG3*, two single guide RNAs (sgRNA) were designed for each gene to create short deletions using the CRISPR P 2.0 online tool (http://crispr.hzau.edu.cn/CRISPR2/) (Liu et al., 2017). The selected sgRNA sequences encompass an *EcoRI* and a *PstI* restriction site for *SlG2* and *SlG3* genes, respectively (Figures S6 and S7). The cloning of the sgRNA sequences was performed as described (Schiml et al., 2016) but using a pDE-Cas9 plasmid providing kanamycin resistance. A pair of primers for each guide was designed, denaturalized and assembled into pENC1.1 (pENTRY) vector previously digested with *BbsI.* The entry vectors contained the corresponding sgRNA expression cassette flanked by *Bsu36I* and *MluI* restriction sites, and by GW recombinant sites to allow both types of interchange with the pDE-Cas9 vector (pDESTINY). The final binary vectors were generated in a two-step cloning process that involved *Bsu36I* and *MluI* digestion-ligation of the first sgRNA into the pDE-Cas9 vector followed by an LR reaction to subclone the second sgRNA of each gene into the pDE-Cas9 vector already containing the first sgRNA.

For activity assays in *E. coli*, full-length *SlG2*, *SlG3*, *slg2-1*, *slg2-2*, *slg3-1* and *slg3-2* sequences were amplified from genomic DNA of the corresponding lines and cloned into the *SmaI* site of the pBluescript SK+ plasmid. All constructs were confirmed by restriction mapping and DNA sequencing. Information about primers used and cloning details are described in Tables S9 and S10, respectively.

### Phylogenetic analyses

Arabidopsis GGPPS and GFPP protein sequences were retrieved from *The Arabidopsis Information Resource* (https://www.arabidopsis.org/) and used as queries to search for tomato homologs on the *Solanaceae Genomics Network* (https://solgenomics.net/) and the *National Center for Biotechnology Information* websites (https://www.ncbi.nlm.nih.gov/) using BLAST. The accession numbers of the identified homologs are listed in Table S1. The presence of transit peptides was predicted using ChloroP and TargetP algorithms (Emanuelsson et al., 2007). Protein and DNA alignments of Arabidopsis and tomato sequences were performed using Clustal Omega (https://www.ebi.ac.uk/Tools/msa/clustalo/) with default settings (Sievers et al., 2011; Sievers and Higgins, 2014). An unrooted phylogenetic tree of Arabidopsis and tomato GGPPS and GFPPS protein sequences was constructed using MEGA6 (Hall, 2013; Tamura et al., 2013). Evolutionary connections among proteins were predicted by the Neighbor-Joining method based on the Poisson model, where pairwise deletion was selected for gap deletions. Bootstrapping was performed on 2,000 pseudoreplicates.

### Subcellular localization assays

Subcellular localization assays were performed by *Agrobacterium tumefaciens*-mediated transient expression in *N. benthamiana leaves* (Sparkes et al., 2006). *A. tumefaciens GV3101* strains were transformed with pGWB405-based constructs (Table S10) and grown on LB plates at 28 °C for 3 days. A single PCR-confirmed colony per construct was grown overnight at 28 °C in 5 mL antibiotic-supplemented LB media and 500 μL of the grown culture were then inoculated in 20 mL of fresh medium. After another overnight incubation, bacterial cells were pelleted and resuspended in infiltration buffer (10 mM MES pH5.5-6, 10 mM MgSO_4_, 150 μM acetosyringone) to a final OD600 of 0.5. To prevent silencing, leaves were co-infiltrated with an *Agrobacterium* strain harboring a HC-Pro silencing suppressor (Goytia et al., 2006). A 1:1 mixture of the two cultures was infiltrated with a syringe in the abaxial part of leaves from 4 to 6-week old *N. benthamiana* plants. Subcellular localization of GFP fusion proteins was determined three days post-infiltration with an Olympus FV 1000 confocal laser-scanning microscope. GFP signal and chlorophyll autofluorescence were detected using an argon laser for excitation (at 488 nm) and a 500–510 nm filter (for GFP) or a 610–700 nm filter (for chlorophyll). All images were acquired using the same confocal parameters.

### GGPPS activity determination

Constructs to produce different truncated GGPPS protein versions were generated in the pET32-GW vector (Table S10). Competent *E. coli Rosetta 2* (DE3) cells (Novagen) were separately transformed with each construct and single transformants were grown overnight at 37 °C in 5 mL of LB medium supplemented with appropriate antibiotics. Then, 250 μL of each overnight culture were diluted in 25 mL 2xYT medium with the required antibiotics and incubated at 37 °C and 250 rpm until reaching an OD600 between 0.5 and 0.8. After inducing the production of the recombinant proteins with 1 mM IPTG, the cultures were grown overnight at 18 °C and 250 rpm. Bacterial cells were harvested by centrifugation at 3,400 rpm for 15 min and pellets were resuspended in 1 mL *Assay* buffer (15 mM MOPSO, 12.5% v/v glycerol, 1 mM ascorbic acid, pH 7.0, 1 mM MgCl_2_, 2 mM DTT). About 0.2 g of zirconium/silica beads 0.1 mm (BioSpec Products) were added and bacterial lysis was carried out in two rounds of shaking for 10 s at a speed of 6.5 in a FastPrep machine (FP120 Bio101 Savant). Cell lysates were subsequently centrifuged for 10 min at 13,000 g and 4 °C, and supernatants were collected for SDS-PAGE and GGPP activity assays. Enzymatic assays were performed in Eppendorf tubes in a final volume of 200 μL containing 25 μL of cell extract, 150 μM IPP and 50 μM DMAPP in *Assay* buffer supplemented with 5 mM Na_3_O_4_V. The reaction mix was incubated for 2 h at 30 °C in mild agitation and stopped by adding 800 μL of 100% methanol / 0.5% formic acid. After vortexing, samples were sonicated for 15 min and centrifuged at maximum speed for 10 min. Supernatants were then evaporated in a SpeedVac concentrator and 80 μL of 100% methanol / 0.65% formic acid were added to the remnant sample. After centrifugation at maximum speed for 15 min, the supernatants were transferred to glass vials. The detection of prenyl diphosphate products by Liquid Chromatography-Mass Spectrometry (LC-MS) was carried out as described (Ruiz-Sola et al., 2016b). Data acquisition and visualization was performed using XcaliburTM software (ThermoFischer Scientific). Kinetic parameters were calculated as detailed (Barja and Rodríguez-Concepción, 2020) using 3 μg of purified SlG1, SlG2, SlG3 and AtG11 enzymes. pET32 constructs were used to produce 6xHis-tagged recombinant enzymes (Table S10) and protein purification from *E. coli Rosetta* cells was carried out using nickel-nitrilotriacetic acid (Ni-NTA) agarose (Qiagen) as described in Barja and Rodríguez-Concepción (2020). Protein concentration was determined according to Bradford method (Bradford, 1976). IPP and DMAPP substrates and FPP and GGPP standards were obtained from Echelon Biosciences. Activity assays in *E. coli* were carried out as described (Beck et al., 2013).

### Gene Co-expression Network (GCN) analyses

GCN analyses were performed as previously described (Ahrazem et al., 2018). Briefly, an ad hoc list of tomato GGPPS and plastidial isoprenoid biosynthetic genes was compiled (Table S2), and used to retrieve all the expression data available for different tomato cultivars/tissues/treatments in the TomExpress database (Maza et al., 2013). Subsequently, pairwise Pearson correlations between each GGPPS gene and each selected isoprenoid biosynthetic input gene was computed for vegetative and fruit tissues and Fisher’s Z-transformation was used to test the statistical significance of pairwise correlations. Finally, pairwise correlations were used to build a heatmap and positive significant correlations (ρ≥0.55) were specifically used to draw gene co-expression networks (Table S3).

### RT-qPCR analyses

Total RNA was isolated from tomato freeze-dried tissue (seedlings, leaves or fruit pericarp) using the Maxwell® RSC Plant RNA Kit with the Maxwell® RSC Instruments (Promega) following the manufacturer’s instructions. RNA was quantified using a NanoDropTM 8000 spectrophotometer (ThermoFischer Scientific) and checked for integrity by agarose gel electrophoresis. The Transcriptor First Strand cDNA Synthesis Kit (Roche) was used to reverse transcribe 0.5 μg of extracted RNA into 20 μL of cDNA, which was subsequently diluted ten-fold and stored at −20 °C for further analysis. Relative mRNA abundance was evaluated via Real-Time Quantitative Polymerase Chain Reaction (RT-qPCR) in a reaction volume of 20 μL containing 10 μL of the LightCycler 480 SYBR Green I Master Mix (Roche), 0.3 μM of each specific forward and reverse primer (Table S9) and 5 μL of cDNA. The RT-qPCR was carried out on a LightCycler 480 Real-Time PCR System (Roche). Three independent biological replicates of each condition and at least two technical replicates of each biological replicate were performed. Normalized transcript abundances were calculated as described previously (Simon, 2003) using tomato *ACT4* (*Solyc04g011500*) or *EXP* (*Solyc07g025390*) as endogenous reference genes. Primer efficiencies were calculated using serial dilutions of genomic or plasmidic DNA. Three biological replicates of cDNA samples from roots of non-mycorrhized and mycorrhized tomato plants (Ruiz-Lozano et al., 2016) were kindly provided by Dr. Juan Antonio López-Ráez.

### Co-immunoprecipitation and immunoblot analyses

Co-immunoprecipitation (Co-IP) assays were performed by transient expression assays in *N. benthamiana leaves* via *A. tumefaciens* as described (Muñoz and Castellano, 2018). Constructs encoding Myc- and HA-tagged tomato GGPPS and PSY proteins (Table S10) were transformed into *A. tumefaciens* GV3101 strains as detailed above. A plasmid containing the Arabidopsis phosphoribulokinase protein with a Myc tag (pGWB417_AtPRK-Myc) was kindly provided by Dr. Ernesto Llamas and used as a negative control. Different *Agrobacterium* infiltration mixtures were prepared and infiltrated into *N. benthamiana* leaves as described (Muñoz and Castellano, 2018). After 3 days, 1.2 g of leaf tissue infiltrated with the same *Agrobacterium* mixtures were frozen in liquid nitrogen and directly stored at −80 °C until use. For crude extracts preparation, frozen leaf samples were ground in liquid nitrogen and incubated in 4 ml lysis buffer (50 mM Tris-HCl pH7.5, 150 mM NaCl, 5% glycerol, 0.05% NP-40, 1 mM MgCl_2_, 0.5 mM PMSF, 1X Sigma protease inhibitor, 10 mM DTT, 2% PVPP) at 4 °C for 15 min using a rotator to form a homogeneous suspension, that was then pre-clarified at 3,000 g for 15 min. Supernatants were cleaned by centrifugation at 16,000 g for 30 min and used for protein quantification (Bradford, 1976). Crude extracts were then adjusted to the same volume and protein concentration with lysis buffer lacking PVPP. An aliquot of each adjusted crude extract was boiled for 10 min in SDS-loading buffer and stored at −20 °C as input sample. 500 μL of each crude extract were incubated overnight with 1 μL of monoclonal αMyc antibody (Sigma) in a rotator at 4 °C. Immunoprecipitation of αMyc interacting protein/complexes was carried out using Pierce Protein A/G Magnetic Beads (ThermoFischer Scientific). After pre-washing the magnetic beads (50 mM Tris-HCl pH7.5, 500 mM NaCl, 5% glycerol, 0.05% NP-40, 1 mM MgCl_2_, 0.5 mM PMSF, 1X Sigma protease inhibitor, 10 mM DTT), the Co-IP sample (crude extract + αMyc antibody) was added and incubated with the beads at room temperature for 1 h with shaking. The beads were then collected with a magnetic stand and repeatedly washed with washing buffer and water. After removing the water from the last washing step, 100 μL of SDS-PAGE loading buffer were added to the beads and boiled for 10 min. Afterwards, the beads were magnetically removed from the supernatants containing the immunoprecipitated complexes and stored at −20 °C. The presence of Myc- and HA-tagged proteins in input and Co-IP samples were detected by immunoblot analyses as described (Pulido et al., 2013) using 1:5000-diluted αMyc (Sigma) and 1:1000-diluted αHA (Roche) as primary antibodies. Horseradish peroxidase (HRP)-conjugated secondary antibodies against mouse and rat IgGs were used in a 1:10000 dilution. WesternBright ECL Western blotting detection kit (Advansta) and Amersham ECL Prime Western Blotting Detection Kit (GE Healthcare) were used for detection and the signal was visualized using the ChemiDoc Touch Imaging System (Bio-Rad).

### Generation of tomato CRISPR-Cas9 mutant plants

*A. tumefaciens* GV3101 strain was used to stably transform tomato MicroTom cotyledons with pDE-Cas9 plasmids harboring two sgRNAs to disrupt *SlG2* and *SlG3* genomic sequences as described (Fernandez et al., 2009). *In vitro* regenerated T1 tomato mutant lines were identified based on kanamycin resistance (100 μg/ml), genotyping PCR and restriction analyses of the amplified regions (Figures S6 and S7). Four T1 lines of each transformation showed biallelic deletions in the designed region. Homozygous T2 lines lacking Cas9 were obtained after segregation. Stable T3 offspring was used for further experiments. For the isolation of double mutants, *slg2-2* and *slg3-1* homozygous plants were manually crossed. Double heterozygous F1 plants were confirmed by PCR genotyping and restriction analyses, and allowed to self-pollinate. The resulting seeds were used to screen the F2 population for double homozygous plants by PCR genotyping and restriction analyses.

### Photosynthetic measurements

Photosynthetic activity was assessed by measuring chlorophyll *a* fluorescence with a MAXI-PAM fluorometer (Walz). The effective quantum yield ϕPSII (ΔF/Fm’) of young and mature tomato leaves was measured as (Fm’−Fs)/Fm’, where Fm’ and Fs are the maximum and the minimum fluorescence of light exposed plants, respectively. The light intensity chosen was 21 PAR (actinic light, AL=2) as previously reported (Llorente et al., 2019). The results are presented as the average of three biological replicates and four different leaf areas for each replicate.

### Metabolite analysis

Carotenoids, chlorophylls and tocopherols were extracted as follows. A mix was prepared in 2 mL Epperdorf tubes with ca. 4 mg of freeze-dried leaf tissue, 375 μL of methanol as extraction solvent and 25 μL of a 10 % (w/v) solution of canthaxanthin (Sigma) in chloroform as internal control. After vortexing the samples for 10 s and lysing the tissue with 4 mm glass beads for 1 min at 30 Hz in the TissueLyser II (Qiagen), 400 μL of Tris-HCl pH:7.5 were added and the samples were again mixed for 1 min in the TissueLyser. Next, 800 μL of chloroform were added and the mixture was again shaken for 1 min in the TissueLyser. Samples were then centrifuged for 5 min at maximum speed at 4 °C. The lower organic phase was placed in a new 1.5 mL tube and evaporated using a SpeedVac. Fruit isoprenoids were extracted using 15 mg of freeze-dried tissue and 1 ml of hexane/acetone/methanol 2:1:1 as extraction solvent. After vortexing and lysing the tissue with the TissueLyser as described for leaves, 100 μL of milli-Q water were added. Then, 1 min of TissueLyser was carried out again and samples were centrifuged for 3 min at 3,000 rpm and 4 °C. The organic phase was transferred to a 1.5 mL tube and the rest was re-extracted by adding 1 mL of hexane/acetone/methanol 2:1:1 solvent, TissueLyser-mixing for 1 min and centrifuging for 5 min at maximum speed and 4 °C. The new organic phase was mixed with that previously extracted and evaporated using the SpeedVac system. Extracted metabolites from leaf and fruit pericarp samples were resuspended in 200 μL of acetone by using an ultrasound bath (Labolan) and filtered with 0.2 μm filters into amber-colored 2 mL glass vials. Separation and detection was next performed using an Agilent 1200 series HPLC system (Agilent Technologies) as previously reported (Fraser et al., 2000). Eluting chlorophylls and carotenoids were monitored using a photodiode array detector whereas tocopherols were identified using a fluorescence detector. Peak areas of chlorophylls (650 nm), carotenoids (470 nm for lycopene, lutein, β-carotene, violaxanthin, neoxanthin and canthaxanthin or 280 nm for phytoene), and tocopherols (330 nm) were determined using the Agilent ChemStation software. Quantification was performed by comparison with commercial standards (Sigma). ABA levels were determined as described (Diretto et al., 2020). Primary metabolites were extracted, annotated and quantified as described previously (Llorente et al., 2019) using approximately 10 mg of lyophilized tissue. Parameters used for peak annotation are provided in Table S6 for leaves and Table S8 for fruits.

## ACKNOWLEDGEMENTS

We greatly thank Juan Antonio López-Ráez for providing cDNA samples of non-mycorrhized and mycorrhized tomato roots; Ernesto Llamas for providing the pGWB417_AtPRK construct, and Albert Ferrer and Laura Gutiérrez for the pDE-Cas9 (with kanamycin resistance) plasmid. The technical support of M. Rosa Rodríguez-Goberna and all CRAG services is also appreciated. This work was funded by the European Regional Development Fund (FEDER) and the Spanish Agencia Estatal de Investigación (grants BIO2017-84041-P and, BIO2017-90877-REDT) and Generalitat de Catalunya (2017SGR-710) to MRC. Support by the collaborative European Union’s Horizon 2020 (EU-H2020) ERA-IB-2 (Industrial Biotechnology) BioProMo project to MRC (PCIN-2015-103), RK and JB (053-80-725) is also acknowledged. CRAG is financially supported by the Severo Ochoa Programme for Centres of Excellence in R&D 2016-2019 (SEV-2015-0533) and the Generalitat de Catalunya CERCA Programme. MVB was funded with a Spanish Ministry of Education, Culture and Sports PhD fellowship (FPU14/05142) and a EU-H2020 COST Action CA15136 (EuroCaroten) short-stay fellowship. ME is supported by a Spanish Agencia Estatal de Investigación (BES-2017-080652) PhD fellowship. IFS is supported by the EU-H2020 Marie S. Curie Action 753301 (Arcatom).

## AUTHOR CONTRIBUTIONS

MVB, ME and MRC designed the research; MVB, ME, GD, IFS, EF, and AF performed research; RK, ARF and JB contributed analytic tools; MVB, ME, GD, IFS, EF, AF, RK, ARF, JB and MRC analyzed data; MVB and MRC wrote the paper.

